# Capturing and analyzing pattern diversity: an example using the melanistic spotted patterns of leopard geckos

**DOI:** 10.1101/2021.03.23.436685

**Authors:** Tilmann Glimm, Maria Kiskowski, Nickolas Moreno, Ylenia Chiari

## Abstract

Animal color patterns are widely studied in ecology, evolution, and through mathematical modeling. Patterns may vary among distinct body parts such as the head, trunk or tail. As large amounts of photographic data is becoming more easily available, there is a growing need for general quantitative methods for capturing and analyzing the full complexity and details of pattern variation. Detailed information on variation in color pattern elements is necessary to understand how patterns are produced and established during development, and which evolutionary forces may constrain such a variation. Here, we develop an approach to capture and analyze variation in melanistic color pattern elements in leopard geckos. We use this data to study the variation among different body parts of leopard geckos and to draw inferences about their development. We compare patterns using 14 different indices such as the ratio of melanistic versus total area, the ellipticity of spots, and the size of spots and use these to define a composite distance between two patterns. Pattern presence/absence among the different body parts indicates a clear pathway of pattern establishment from the head to the back legs. Together with weak within-individual correlation between leg patterns and main body patterns, this suggests that pattern establishment in the head and tail may be independent from the rest of the body. We found that patterns vary greatest in size and density of the spots among body parts and individuals, but little in their average shapes. We also found a correlation between the melanistic patterns of the two front legs, as well as the two back legs, and also between the head, tail and trunk, especially for the density and size of the spots, but not their shape or inter-spot distance. Our data collection and analysis approach can be applied to other organisms to study variation in color patterns between body parts and to address questions on pattern formation and establishment in animals.

## Introduction

Animal color patterns vary within and among individuals, including variation among distinct body parts such as the head, trunk, tail, wings, or ventral or dorsal sides, possibly in response to different selection pressures (Caro, 2005; Forsman et al., 2008; Allen et al., 2020). Color patterns may differ in qualitatively obvious ways, such as stripes on the tail and spots on other parts of the body, or in more subtle ways, such as spots of different density or sizes (e.g., Figure 1).

**Figure 1:**
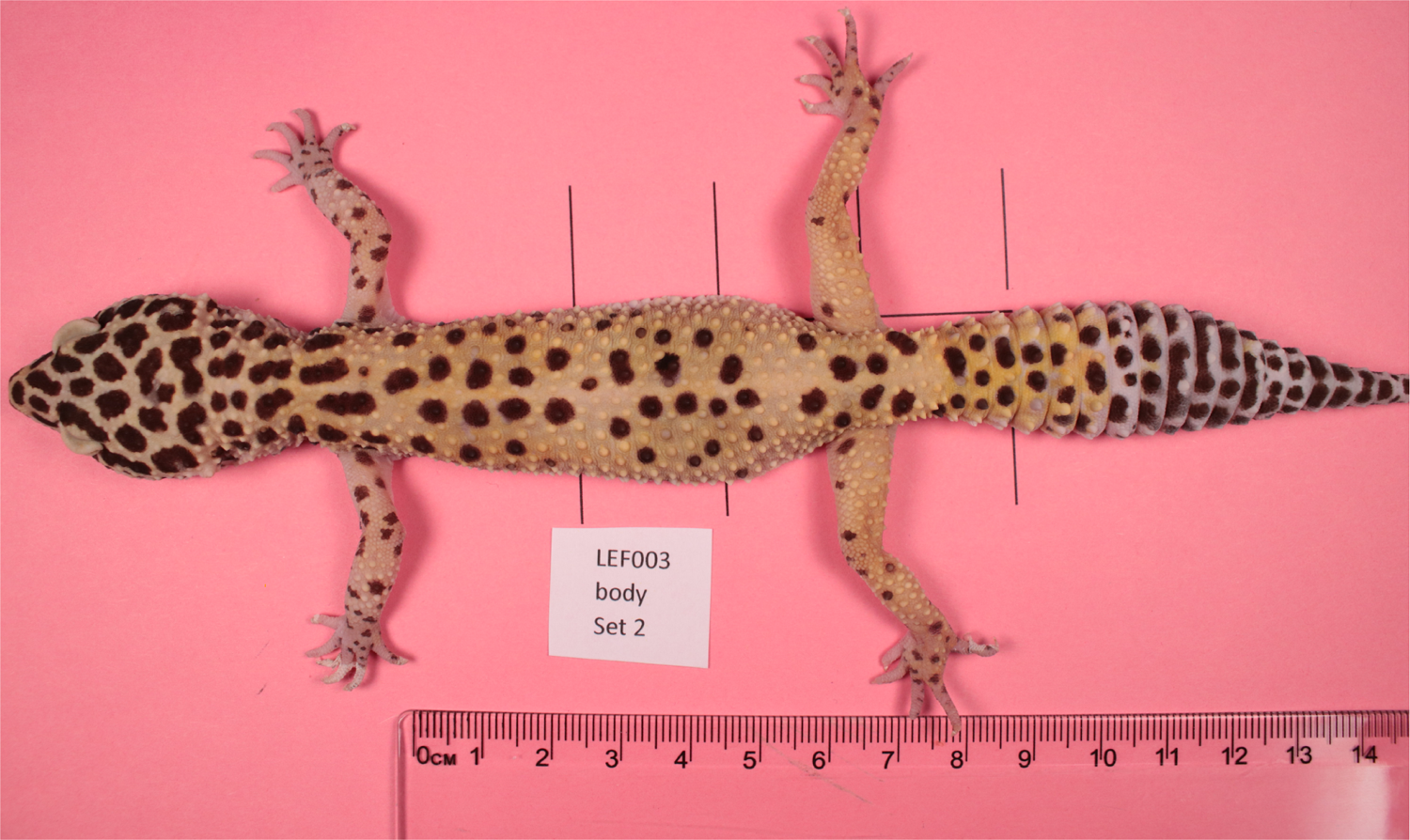
Adult leopard gecko (animal #21003). Note the differences in patterning on the head, trunk, legs and tail.

Variation in color pattern is considered a classical example of an adaptive trait, as it is often involved in communication among conspecifics, intrasexual competition, and antipredator functions (Caro, 2005; Gomez et al. 2007, Tibbetts and Dale, 2004, Solan et al. 2019).

Although color patterns have been studied extensively because they are involved in many functions essential to the survival and reproduction of organisms, describing and quantifying pattern variation in a multivariate manner is still challenging. Color patterns are generally described in terms of macroscopic differences, such as spots, stripes or labyrinthine organization (e.g., Miyazava et al. 2010, Allen et al. 2020, Kuriyama et al. 2020), with pattern variation for the same pattern type often quantified using landmarks obtained on homologous pattern features (e.g., van Belleghem et al 2018, Bainbridge et al. 2020, Prinsloo et al. 2020) or by focusing on differences in pattern elements, coarsely defined in terms of relative size and position (e.g., van den Berg et al. 2020 and references therein). However, complex patterns with high degree of dissimilarity within and among individuals in terms of shape, clustering, size and position of the pattern elements may require the development of new methods to finely capture these differences (see for example Lee et al. 2018 and references therein) This is particularly pertinent if the shape and density are irregular and do not fit within specific pattern categories, such as stripes or spots (Solan et al., 2019; Troscianko et al., 2017; Lee et al., 2018; Miyazawa et al. 2010; McGuirl et al., 2020; Allen et al., 2020).

Color pattern can be studied using pattern recognition, which broadly speaking deals with classification of image patterns through extraction of significant features (Zerdoumi et al., 2018). Methods in this area generally consist of machine learning techniques, that is, the prediction systems based on an existing data set. For example, this could entail classification of skin patterns based on a large training data set of skin pattern images. While this approach is certainly viable for synthetic data, i.e. computer-generated patterns (McGuirl et al, 2020), where data with thousands of patterns can be amassed easily, this approach may not be feasible for actual images of live animals, where the process of image acquisition is laborious and time consuming. In addition, the use of machine learning techniques would not necessarily provide qualitative insights into what would make two or more patterns similar or different (Domingos, 2012; Zerdoumi et al., 2018), therefore impeding investigation of what elements of the pattern for example may be more or less variable, constrained or under selection.

In this article, we address the problem of describing and quantifying variation in melanistic color patterns in live geckos via computing fourteen different indices, such as the fraction of dark areas to light ones, or the mean size, number and shape of pattern features. Each of these different indices captures only one aspect of the pattern, but collectively, they yield a comprehensive characterization of the pattern itself. Thus, each pattern of the seven body parts studied for each individual corresponds to a point in an abstract 14-dimensional space (here called “pattern space” or “phenotype space”). Our approach is similar to that of Lee et al. (2018) who used 11 indices to characterize giraffe coat patterns and Miyazawa et al. (2010), who used two indices to describe salmonid fish skin patterns. In contrast to those papers, however, we not only compare single indices between individuals and among individuals, but also consider different ways to measure the overall similarity of two patterns based on their distance in pattern space, taking into account biological information in the data set. In this, our approach is therefore innovative. Arguably, there is no canonical distance function on this multidimensional space to measure overall similarity of patterns, and so we employ two different notions of a metric, both weighted Euclidean distances. These two distances differ in the type of biological information that they may provide. The first distance, the standard Mahalonobis distance, essentially weights each principal component by the inverse of its variance (Mahalanobis, 1927; Krzanowski, 2000). This standard metric weighs all data points equally and thus does not take into account any inherent structure of the data set, as for example any developmental relationship among body parts. The second distance that we selected is instead a measure that weights differences in patterns by the influence of random noise in the developmental process. We call this distance the “Developmental Noise distance”. In it, differences in indices for which developmental noise has a small contribution are weighted heavier than differences in indices for which it has a larger contribution. The use of these two distance measures therefore not only permits to quantitatively describe and statistically test pattern variation, but also to help understanding the developmental sources - genotypic, environmental, or stochastic - of this variation.

We apply our new approach to capture pattern data and our pattern distance measures to investigate the variation of melanistic skin patterns among distinct body parts for 25 leopard geckos (*Eublepharis macularius*) (Figures 1 and 2 and Table 1). We use these data to analyze different pattern indices and calculate their correlation to infer the order of pattern formation and establishment across the body and to characterize pattern variation on the different body parts within and among geckos. Finally, by comparing the within-individual left-right variation in leg patterns, which is likely due to developmental noise, to the between-individual variation, which is due to genetic and environmental differences in addition to developmental noise, we quantify the influence of developmental noise on pattern variation for the leopard gecko.

**Figure 2:**
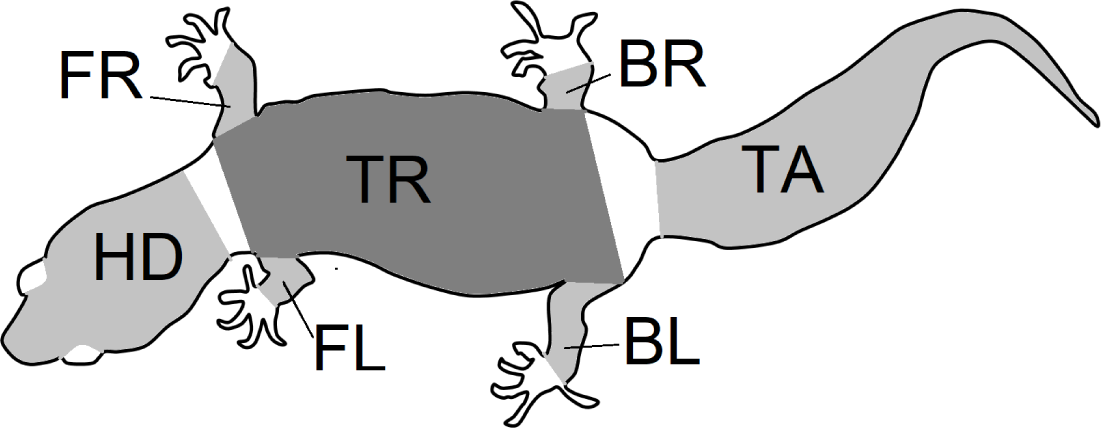
Outline of the gecko body showing the seven regions of patterns that were isolated from gecko images. Each region was photographed separately, and the limbs were gently stretched during photographing.

**Table 1:** Gecko morph (“normal” or “lemon frost” - LF), source, ID, sex (M=male, F=female), and presence of a visible (YES) or not discernible (NO) melanistic pattern on the seven different body parts for each gecko. “BL” and “BR” indicate the back right and left legs, respectively; “FL” and “FR” indicate the frontal left and frontal right legs, respectively. “TR”, “TA” and “HD” indicate the dorsal part of the trunk, tail and head of the animal, respectively. (Sources of obtained geckos: w—Greg Watkins—Colwell, g—Tony Gamble, b—Backwater Reptiles, t—Ron Tremper)

|  | Morph | Source | ID | Sex | BL | BR | FL | FR | TR | HD | TA |
| --- | --- | --- | --- | --- | --- | --- | --- | --- | --- | --- | --- |
| 1 | Normal | w | 10001 | F | YES | YES | YES | YES | YES | YES | YES |
| 2 | Normal | g | 21001 | F | YES | YES | YES | YES | YES | YES | YES |
| 3 | Normal | g | 21002 | F | YES | YES | YES | YES | YES | YES | YES |
| 4 | Normal | g | 21003 | F | YES | YES | YES | YES | YES | YES | YES |
| 5 | Normal | g | 22010 | M | YES | YES | YES | YES | YES | YES | YES |
| 6 | Normal | b | 22013 | M | YES | YES | YES | YES | YES | YES | YES |
| 7 | Normal | b | 41002 | F | YES | YES | YES | YES | YES | YES | YES |
| 8 | Normal | b | 41004 | F | YES | YES | YES | YES | YES | YES | YES |
| 9 | Normal | b | 42001 | M | YES | YES | YES | YES | YES | YES | YES |
| 10 | Normal | b | 42002 | M | YES | YES | YES | YES | YES | YES | YES |
| 11 | Normal | b | 42003 | M | YES | YES | YES | YES | YES | YES | YES |
| 12 | Normal | b | 42004 | M | YES | YES | YES | YES | YES | YES | YES |
| 13 | Normal | g | 22011 | M | NO | NO | YES | YES | YES | YES | YES |
| 14 | Normal | b | 22012 | M | NO | NO | YES | YES | YES | YES | YES |
| 15 | Normal | w | 10002 | F | NO | NO | NO | NO | YES | YES | YES |
| 16 | Normal | g | 21008 | F | NO | NO | NO | NO | YES | YES | YES |
| 17 | Normal | b | 41001 | F | NO | NO | NO | NO | YES | YES | YES |
| 18 | Normal | b | 42005 | M | NO | NO | NO | NO | YES | YES | YES |
| 19 | Normal | g | 21007 | F | NO | NO | NO | NO | NO | YES | YES |
| 20 | Normal | b | 41003 | F | NO | NO | NO | NO | NO | YES | YES |
| 21 | LF | t | 21019 | F | YES | YES | NO | NO | YES | YES | YES |
| 22 | LF | t | 21020 | F | YES | YES | NO | NO | YES | YES | YES |
| 23 | LF | t | 22016 | M | YES | YES | NO | NO | YES | YES | YES |
| 24 | LF | t | 22014 | M | YES | NO | NO | NO | YES | YES | YES |
| 25 | LF | t | 21018 | F | NO | NO | NO | NO | YES | YES | YES |

Among vertebrates, lizards have often been used as ideal models to study the evolution of color and color pattern in relationship to other ecological, biological, and behavioral traits (e.g., Olsson et al., 2013; Pérez I de Lanuza and Font, 2016; Murali et al., 2018; Allen et al., 2020).

Specifically, the leopard gecko is an ideal organism on which to study pattern development (e.g., Chang et al., 2009). This species is commonly bred in captivity to obtain distinct colors and color patterns, a major advantage when trying to unveil the mechanisms producing variation at these traits (Cieslak et al. 2011). Furthermore, our previous work on color patterns on the head of this species (Kiskowski et al., 2019) suggests that developmental noise may be an important contributor to its variation.

This work therefore not only proposes a novel approach to analyze similarities in pattern space within and among individuals, but also contributes to our understanding of melanistic pattern formation and establishment in the leopard gecko. The data capture and analysis methods presented here can also be applied to study variation in color pattern elements for developmental, ecological and evolutionary purposes in other organisms. Furthermore, our previous work used mathematical modeling of the process of skin pattern formation to elucidate the influence of developmental noise on patterning (Kiskowski et al., 2019). Many different other mathematical models for skin patterning have been proposed (e.g. Murray, 2002; Cruywagen et al., 1992; Painter, 2001; Cooper et al., 2018; Kondo et al., 2009). In this context, developmental noise can be modeled for example by using random initial conditions for the equations governing pattern formation. Because of this randomness, the resulting pattern is not deterministic, but rather a certain range of possible patterns - all depending on randomization of these initial conditions - may be generated with an associated probability density function. This range can be thought of as a set of points in the 14-dimensional pattern space provided by the measures used in the current work. This approach and the empirical data can then be used to assess the validity of the models, and thus in turn biological insights (see McGuirl et al. 2020 for an example) into the patterning process.

Nevertheless, depending on the gecko configuration, viewing angles and the region shapes were irregular. An image processing algorithm was used to identify the pattern that was viewable within each region irrespective of the shape. The abbreviations BL (left back leg), BR (right back leg), FL (left front leg), FR (right front leg), HD (head), TR (trunk) and TA (tail) are as in Table 1 and are used throughout this article.

## Methods

### Ethical statement

All experiments were carried out in accordance with George Mason University animal use (IACUC) protocol # 1430668.

### Geckos and Photographs

For this study we used a total of 25 live adults of *Eublepharis macularius*, the leopard gecko, giving a total of 132 patterns on various body parts. 20 geckos had an overall “normal” pattern morphotype (melanistic - black - spots on a yellowish/brownish background), while five geckos had a “lemon frost” morphotype with melanistic patterns (Szydłowski et al. 2020, Guo et al. 2020; Figure A1 in the Appendix for full body images of all geckos, Table 1 and Supplementary Material for details on the origin of the geckos).

We photographed the geckos one at the time by placing each one of them on a smooth surface covered with colored paper (RGB=[100,60,65], Figure 1) chosen to contrast well with the full set of geckos in at least one color channel) (Figure A1 in the Appendix). Perpendicular reference lines were printed on the colored paper to ensure placement of the gecko in the same position across picture sets. For each gecko, we obtained four picture sets to measure the error associated with the data capture (See Supplementary Material for further details).

### Image Analysis

The melanistic spotted patterns were studied on the head, four limbs, dorsal trunk, and tail of each of the 25 geckos. There were thus 25×7=175 separate body parts analyzed in this work (Figure 3). Not all these body parts showed melanistic spotted skin patterns and only twelve of the geckos had qualifying spotted patterns on all seven body parts (Table 1; see details below for how qualifying patterns were recognized).

**Figure 3:**
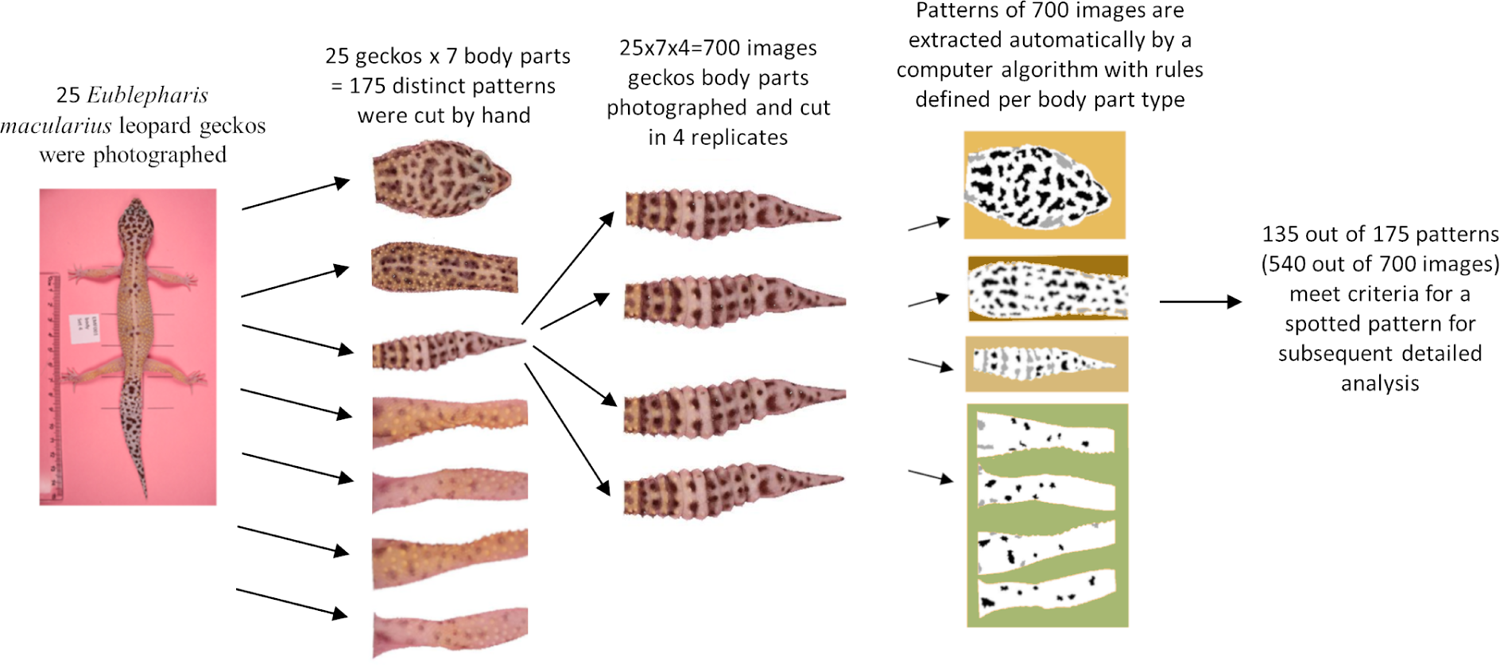
Flow chart of image acquisition and processing.

For our analyses, each of the seven regions for each gecko was isolated by automated removal of background pixels from the images where possible and additionally by hand using the GIMP photo editing tool (Figure 3). When cutting the regions, we worked along the natural boundary of the body and had defined rules for the edges of the body region (e.g., the trunk was separated from the head by a straight line segment connecting the two most anterior points where each front legs met the main body, and the legs were separated from the body using a straight line segment perpendicular to the limb that was the most proximal line segment that could be drawn without including any portion of the trunk. See Figure A3 for more details). For each of the 700 images (25 geckos, 7 body parts, 4 independent photos of each), we identified and isolated the spotted melanistic pattern as a simple binary pattern of black pixels on a white background (i.e., in every image, each pixel has either value of 1 = black or 0 = white). Using criteria to help in isolating the melanistic spotted pattern amongst other patterns of the skin and background noise of the image, a threshold was applied to each of the regions to define the binary pattern of black spots on a white background (see below for details and Table 2). A spot identification algorithm was applied for each type of body pattern to identify spots. A final image processing step with Matlab removed stray pixels, filled in holes, and smoothed the contour of the spots to generate the final spot patterns used for measuring the pattern statistics. The Matlab code for these scripts are available in the Supplementary Material (after acceptance of the manuscript for publication).

**Table 2:** Algorithm criteria for determining the threshold for spots for the four body parts (limbs, head, trunk, and tail) including the color channel that was used for identifying spots (the green channel, for all) and the color channel that was used for identifying regions of the picture requiring lighting adjustment (the green channel for the legs and the blue channel for the trunk; the mean and standard deviation of the pixel values in the region are denoted as µ_G_, σ_G_ and µ_B_, σ_B_, respectively) and the final threshold rule that was applied to the green channel.

| Body Part Type | Channel Used to Compute the Mean Pixel Intensity $\mu$ | Channel and threshold Used to Identify Shadows | Threshold |
| --- | --- | --- | --- |
| Limbs | Green | shadows: $\text{green} < \mu_G - 0.3\sigma_G$<br>glare: $\text{green} > \mu_G + 0.3\sigma_G$ | maximum of these two quantities:<br>$\mu_G - 0.85\sigma_G, 60$ |
| Head | Green | N/A | maximum of these two quantities:<br>$\mu_G - 0.50\sigma_G, 60$ |
| Trunk | Green | shadows: $\text{blue} < \mu_B - 0.85\sigma_B$ | maximum of these two quantities:<br>$\mu_G - 0.85\sigma_G, 60$<br>additionally, maximally: 108 |
| Tail | Green | N/A | maximum of these two quantities:<br>$\mu_G - 0.85\sigma_G, 60$ |

#### Limb, Trunk, Head, and Tail Spot Identification Algorithms

Due to morphological differences in the four body parts, the algorithms used to determine the melanistic pattern from the photographic images varied for each body part, but was otherwise applied the same way to every gecko image for uniformity and consistency. In a preliminary step that was completed by trial and error and evaluated by eye, all rules for the algorithm (for example, which color channel to use, whether the lighting across the images would be adjusted, and the threshold darkness criterion for which a pixel would be identified as melanistic or not) were chosen for each body part for a good fit between paucity (to minimize the number of criteria and minimize differences between the four algorithms) and robust ability (as determined by eye) to capture the melanistic patterning for the greatest number of gecko images.

All of the differences between the four algorithms are summarized in the Supplementary Material. See Supplementary Material for images of all 7 body parts and binarized images for 25 geckos (times 4 repeated measurements); also see Figure A2 in the Appendix for a representative example of the binarized images for one body part for one gecko.

### Image Length Scale

For each image, the length scaling factor (length per pixel) was computed via determining the number of pixels for one centimeter using the imaging software GIMP. This was used in converting values measured in pixels to lengths, see Table 3.

**Table 3:** Indices used in this work to describe patterns. Like Miyazawa et al. (2010), we include a measure for the ratio of black to white pixels (FM) and a measure of the circularity of spots (EE) among the 14 variables used in this work. See Supplementary Material for definition of interior spots.

| *Abbr.* | *Name* | *Units* | *Description* | *Spot type used for the computation* |
| --- | --- | --- | --- | --- |
| FM | fraction of melanistic area | dimensionless | Ratio of black pixels to all pixels in binarized image (ratio of melanistic area to total area) | All Spots |
| SS | mean spot diameter ('spot size') | cm | Mean of the length of the major axes of the ellipses that have the same second moment as the spots (Matlab function *MajorAxisLength*) | Interior Spots |
| SSD | st.dev. of SS | cm | Standard deviation of the lengths of major axes used in definition of SS | Interior Spots |
| EE | mean ellipticity | dimensionless | Mean ratio of major axes to minor axes of ellipses that have the same second moment as the spots (Matlab function *Eccentricity*) | Interior Spots |
| EED | st. dev. of EE | dimensionless | Standard deviation of the ratios used in definition of EE | Interior Spots |
| PL | peak length | cm | Measure of characteristic wavelength; (typical distance between spots; see Miura et al. (2000)) | All Spots |
| MD | mean minimum distance | cm | Mean of the mean distance of the centroid of a spot to the closest three other spots' centroids | All Spots |
| MDD | minimum distance st. dev. | cm | Standard deviation of the distances used in the definition of MD | All Spots |
| SA | spot area | cm<sup>2</sup> | Mean area of spots | Interior Spots |
| SAD | st. dev. of spot area | cm<sup>2</sup> | Standard deviation of the spot areas | Interior Spots |
| SI | spot intensity | dimensionless | Mean green values (G) of the RGB values of spots (between 0 and 255 for each pixel) | All Spots |
| SID | st. dev. of spot intensity | dimensionless | Standard deviation of the intensities used in the definition of SI | All Spots |
| EL | mean spot elongation | dimensionless | Mean of the spot elongation $EL = \frac{area}{2d^2}$ where $d$ is the thickness of the spot (number of erosion steps needed before the pot disappears) | Interior Spots |
| ELD | st. dev. of spot elongation | dimensionless | Standard deviation of the spot elongation | Interior Spots |

#### Application of a Threshold to Define Spots

Each pixel of an image has an R, a G and a B value, which are integers that each range from 0 to 255. We used the green channel G as a measure of the intensity of a spot (see Table 2). Threshold intensities for determining whether a pixel in an image was dark enough to be a pixel within a melanistic spot was based on the average intensityµ and the standard deviation σof the intensity within the region. For the legs, trunk and tail, a threshold of µ − 0. 85σ was determined by eye to best capture the melanistic pattern across the entire set of geckos. For the head, which had a larger fractional area of melanistic spots, a threshold µ − 0. 5σwas determined to identify the set of melanistic spot pixels best. A high fraction of melanistic area relative to total skin area (combined with exceptionally dark spots) would decrease the average intensity to values that were too low to identify more lightly colored spots so a minimum was applied that the threshold for the value of a pixel would be at least 60 out of 255. Pixels with values this low were invariably likely to be melanistic. A maximal threshold of 108 out of 255 was applied to the trunk since a very low fractional melanistic area could cause the average intensity to be too large for good spot detection; however, higher threshold values were frequently appropriate for other body parts (for the legs for example) so no maximal threshold was applied for parts other than the trunk.

#### Final Pattern Processing

The application of the threshold identified a set of pixels that are darker than the threshold value for each evaluated image. This set of pixels included stray pixels as well as larger contiguous areas of pixels that were likely to be pigmented spots. A minimum spot size of 350 pixels (∼0.5 mm^2^) was required for the spot to be qualified as such in this work. This number was determined by visual inspection to remove stray pixels and very small dark artifacts. Although the size of pigmented regions varied (especially the size of these regions could be very large on the head and the trunk), none of the pigmented areas that should be identified as spots were smaller than about 750 pixels in total area, so the minimum spot size was chosen to be smaller than the pigmented areas the algorithm needed to identify but larger than most artifacts.

In the last processing steps, any holes within pigmented regions (see Supplementary Material Figure S1, Panel F) were filled, and the contour of the spots was smoothed using successive dilations and erosions (this removes small scale granular effects at the edges of spots without changing the shape of the spot). See the Supplementary Material for details.

#### Final Pattern Classification

For both the limb and trunk patterns, the spotted pigmented pattern would occasionally be very faint and barely present or not present at all. To distinguish among these, an image was classified as a patterned only if there were at least four interior spots for limb patterns and at least six interior spots for the trunk and tail. The head was invariably well-patterned, with at least 26 spots found on the heads of all the geckos, so no minimum was applied.

### Description of indices

For each of the 25 geckos, we excluded images without patterns as determined by the algorithm described above (Tables 1 and 2) giving a total of 132 qualifying patterns (14-16 patterns for each of the 4 legs, 23 trunk patterns, 25 head patterns and 25 tail patterns). For each such qualifying combination of body parts and gecko, we used Matlab to calculate the 14 indices summarized in Table 3. The value of each index is the average of the 4 independent measurements, giving a total of 132×4 =528 images that were analyzed. For an assessment of the measurement error, see below and the Supplementary Material.

### Definition of distances on pattern space: Mahalanobis and Developmental Noise distances

Each qualifying pattern (one of 7 body parts of one of 25 geckos) is described by the 14 indices in Table 3. Thus we can think of a pattern as a point x = _(_x_1_, …, x_14)_^T^ in a 14-dimensional pattern space. (The superscript “T” denotes the transpose). To quantify the similarity of two patterns, we measure the distance between two points in this pattern space. We consider two different definitions of a distance (metric) on pattern space. Let x = _(_x_1_, …, x_14)_^T^ and y = _(_y_1_, …, y_14)_^T^ denote two points. The first distance we consider is the standard Mahalanobis distance given by

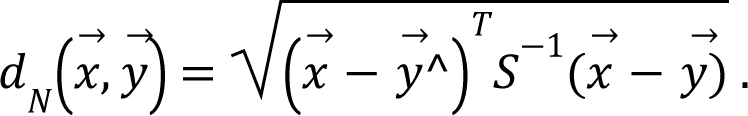

Here S is the covariance matrix of the complete data set. One can think of the Mahalanobis distance as the Euclidean distance computed after transforming the data to principal components and normalizing each principal component (Krzanowski, 2000). The advantage of this method over the Euclidean distance on the untransformed pattern space is that the principal components are uncorrelated, and so the Mahalanobis distance is not skewed by correlations between the different indices. It is generally regarded as an appropriate generic choice for a statistical distance in sample spaces with differential variances and correlations (Krzanowski, 2000). As a generic choice however, the Mahalanobis distance does not take into account any specific information from the particular structure of our data set. In the case at hand, the data points can be grouped by animal, by body part, or for instance by pairs of front legs or back legs of the same animal. In general, differences among patterns can be attributed to differences in the genotype, the environment experienced, and developmental noise. The differences among patterns within the pairs of data points describing the front legs or the back legs of the same animal are likely primarily the result of developmental noise, as opposed to two different genotypes or different environmental conditions. Data obtained from the legs allow then to separate the influence of developmental noise from genetic and environmental factors. We take this into account and define a second metric called “Developmental Noise metric”, defined as a weighted Euclidean metric:

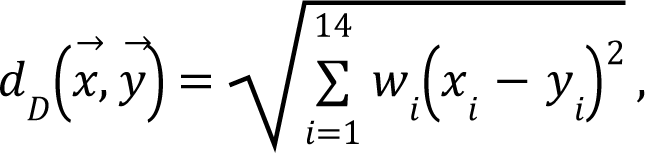

where the weights w_i_, i = 1, …, 14 are defined via the standard deviation of the two front leg i patterns of each gecko. More specifically, for the index i (with 1≤i≤14), let S^n^_i_ denote the variance of the indices of the two front leg patterns of the nth gecko (we only included geckos that have patterns on all four legs; see Table 1). Then we define the weight w as the inverse of the mean of the variances S^n^_i_, i.e.

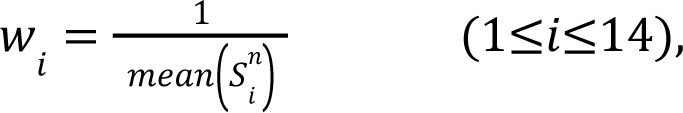

where the mean is taken over all geckos who have patterns on front legs. Note that this distance function is scale invariant, i.e. independent of the units used for the different indices. The reason for using these weights is that the mean variance between the two front legs is a rough measure for the importance of noise in the establishment of the pattern; thus effectively the fewer influence noise has on a measurement, the more weight it is given in the computation of the distance between patterns. While this distance takes into account the special structure of the data set, its computation is based only on a subset of the data, namely leg patterns of those geckos that have patterned front legs. This is in contrast to the standard Mahalanobis metric, which ignores the special structure, but is based on all data points. It is a priori not clear which of these distances is more appropriate, and for this reason we use both in the following analyses. In fact, we found that in general, the results for these two metrics agree qualitatively, giving added confidence in our results (see Results section). Many other reasonable concepts of distances on pattern space are possible. Results are reported in all cases for the *squares* of the distances.

### Quantification of measurement error

We took four independent photos of each body part, where the animal was picked up and rearranged for each repetition so that the four measurements would be independent. A measurement error was introduced by slight differences in the rotation and placement, especially for the limbs and tail. To quantify the measurement error, we took two approaches: in the first, we compared the mean distances in pattern space between the different measurements to the mean between-individual distances of the same body part. The second consisted of a two-way ANOVA test for the front legs and the back legs, where the two factors are “sides” (*S*, fixed) and “individuals” (*I*, random). For more details, see the “Results” section below.

### Statistical Analysis and Software

Images of various body parts were extracted via automated removal of background pixels and consequent manual selection from photos of the geckos with the image editor GIMP. All computations for image analysis and statistical analysis were performed with Matlab. The computations of statistical significance of results (p-values) were performed either with standard statistical tests as implemented in Matlab where indicated, or via nonparametric permutation tests (see the Appendix for details on the procedure). The Matlab code is available in the Supplementary Material (after acceptance of the manuscript for publication).

## Results

### Patterning

Table 1 shows that not all geckos had melanistic patterns that were identified on all body parts. In some cases there was no visible spot pattern by eye as well, in some cases the pigmented melanistic pattern visible by eye was very light and not discerned by the algorithm.To ensure a robust pattern with enough spots to measure average characteristics, we included a spot pattern only if the algorithm identified a minimum number of spots (at least 4 or 6 interior spots). All of the geckos had patterning on the heads and tails, only two were missing patterns on the trunk, and approximately half (13/25) of the geckos were missing patterns on their legs. There is furthermore an identifiable hierarchy of patterning {head;tail}→ {trunk}→ {front legs;back legs} for each individual gecko, where absence of patterns in one of the body parts entails absence of patterning in all “downstream” body parts with respect to this hierarchy. For instance, absence of patterning on the trunk means that all legs have no patterns as well. There was no clear hierarchy between the front legs and back legs. Of those missing patterns on legs, most geckos (7/13) were missing patterns on both sets of legs, 2 were missing patterns on their back legs only, and 4 were missing patterns on their front legs only. Although our sample size is limited for the “lemon frost” morph, there appeared to be an effect of morphotype: for the “normal” morphotype, absence of patterns on the front legs always meant that the back legs were also unpatterned, whereas this was reversed for this morph, since no gecko had patterned front legs.

### Measurement error

We took four independent photos of each body part. To estimate the amount of measurement error in each of the 14 indices, we took two different approaches (see Methods).

In the first, we consider body part patterns as points in 14-dimensional phenotype space and measure the distances between them. We determined first the mean distance of the four repeated measurements of the same body part of the same animal from their centroid. This is an absolute measure of the measurement error. Table A1 in the Appendix lists these errors for all seven body parts as well as the relative error as the ratio of these errors relative to i) the mean *between-individual* distances for the same body part, or ii) the corresponding *within-individual* distances for front or back legs (comparing the left and right leg of the same gecko). The results for both the Mahalanobis distance and the Developmental Noise distance are listed. In all cases, the mean within- or between-individual distances were significantly greater than the mean distance due to the measurement error, with factors varying from 2.2 (back leg, within-individual distance relative to measurement error, Mahalanobis distance) to 86.1 (tail, between-individual distance relative to measurement error, Developmental Noise distance). A factor of 1 would mean that within- or between-individual distances were of the same magnitude (and thus indistinguishable) from measurement error. However these distances were at least twice as large (and often much larger), indicating that the measurement error is relatively small.

In the second approach for characterizing the measurement error, we conducted a two-way ANOVA test where the two factors are “sides” (S; fixed) and “individuals” (I; random) separately for both pairs of front legs and pairs of back legs for each of the 14 indices. This is a standard approach to compare the relative contributions of nondirectional asymmetry (biological) and measurement error (technical variation) in the investigation of paired structures (Palmer and Strobeck, 1986; Merila and Biorklund, 1995; Breuker et al., 2006). For each index, an F-test yielded that nondirectional asymmetry is making a significant contribution to the variation observed relative to measurement error. The F-values had a median value of 6.8, meaning that the measurement error made up about 15% (median value) of the total observed variation between the left and right leg patterns. See Table A2 in the Appendix for details.

### General pattern variation across geckos and body parts

For each body part of each of the 25 studied animals, we determined the value of each of the 14 indices listed in Table 3 via the mean of four repeated independent measurements.

An examination of the coefficient of variation (ratio of standard deviation and mean) for the 14 indices shows that melanistic pattern is highly variable for measures that concern the proportion of melanistic areas (FM), how large these individual areas are (SA and SSD), and, to a lesser extent, what their typical distances are from each other (PL and MD) (Table A4in the Appendix).

This is not an artifact of the measurement error, which actually tended to be larger for MD than the other indices by some measures (see Table A2). In essence, spots can be in higher or lower density across geckos and body parts and larger or smaller. However, once a melanistic pattern is established, the spots are all similar in shape (EE, EL). Table 4 displays the pairwise Pearson correlation coefficients showing how much one index is correlated with another. For example, measures of the size of spots such as FM, SS and SA are strongly positively correlated. The two indices of the typical wavelength (roughly representing the typical distance among melanistic areas), PL and MD, are also positively correlated. It is also noteworthy that EE, which quantifies aspects of the shape of individual spots, is only weakly correlated with the other indices, with the exception of the mean elongation (EL), which also quantifies aspects of the shape of individual spots. This indicates that the spot shape only weakly depends on size or distribution of the spots. A negative correlation value indicates that two indices are anti-correlated. For example, fractional melanistic area (FM) and peak length (PL) are moderately anti-correlated since fractional melanistic area – the proportion of melanistic area to total skin area-tends to increase with the number of spots while peak length a measure of the typical distance between spots - decreases with the number of spots. EE and EED are moderately anti-correlated, which indicates that spots with large eccentricity tend to have a lower variation in their eccentricity, which indicates an eccentricity that is non-random.

**Table 4:**
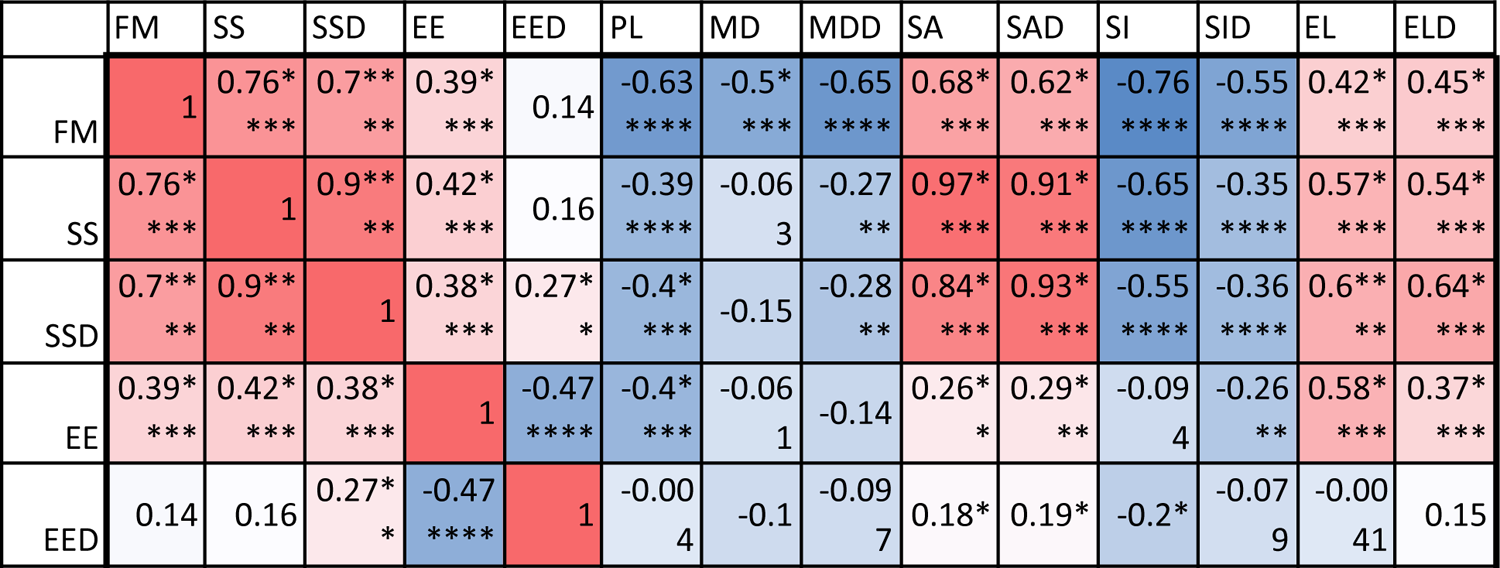
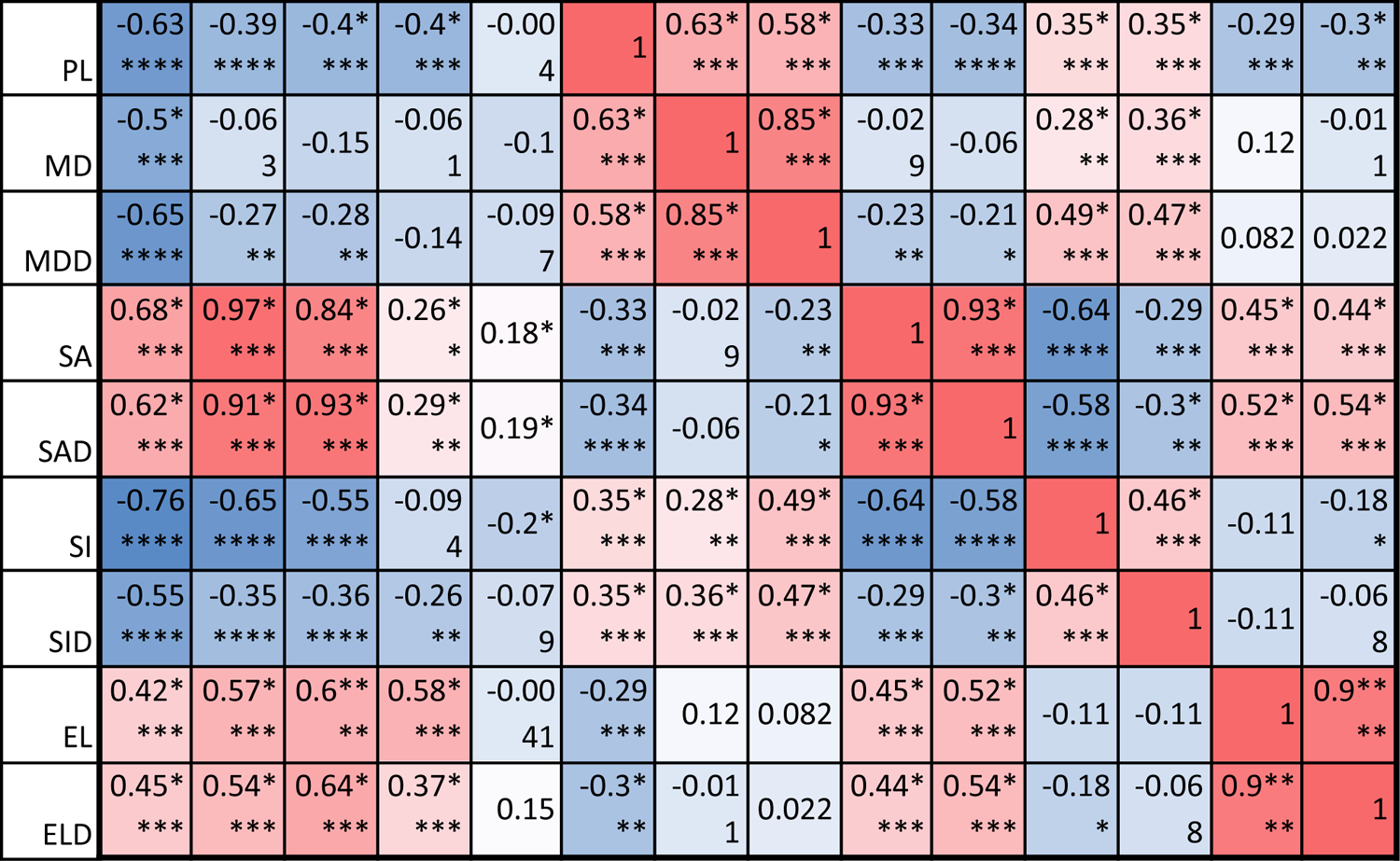
Correlation matrix with values color coded from negative (blue) to positive (red). Symbols for each of the 14 indices are as in Table 3. One star (*) indicates p-values less than 0.05, ** p-values less than 0.01, *** p-values less than 0.001, **** p-values less than 0.00001. Data obtained on all the 25 geckos together independently of morphotype. See the Supplementary Material for differences among morphotypes.

We also computed the correlation coefficients of within-individual indices of the various body parts. The results are summarized in Table 5. A total of 127 out of the 294 correlation coefficients were statistically significant at the 0.05 level of significance (39.8%), meaning that we can statistically reject the hypothesis that variation of these indices is independent among these body parts. The number of statistically significant coefficients varied substantially by the pair of body parts. Correlation is significant for most indices for the two front leg patterns and the head and tail patterns (FL-FR, HD-TA; each 11 out of 14). To a somewhat lesser extent, the indices for the two back legs tended to be correlated (BL-BR; 9 out of 14 significant).

**Table 5:**
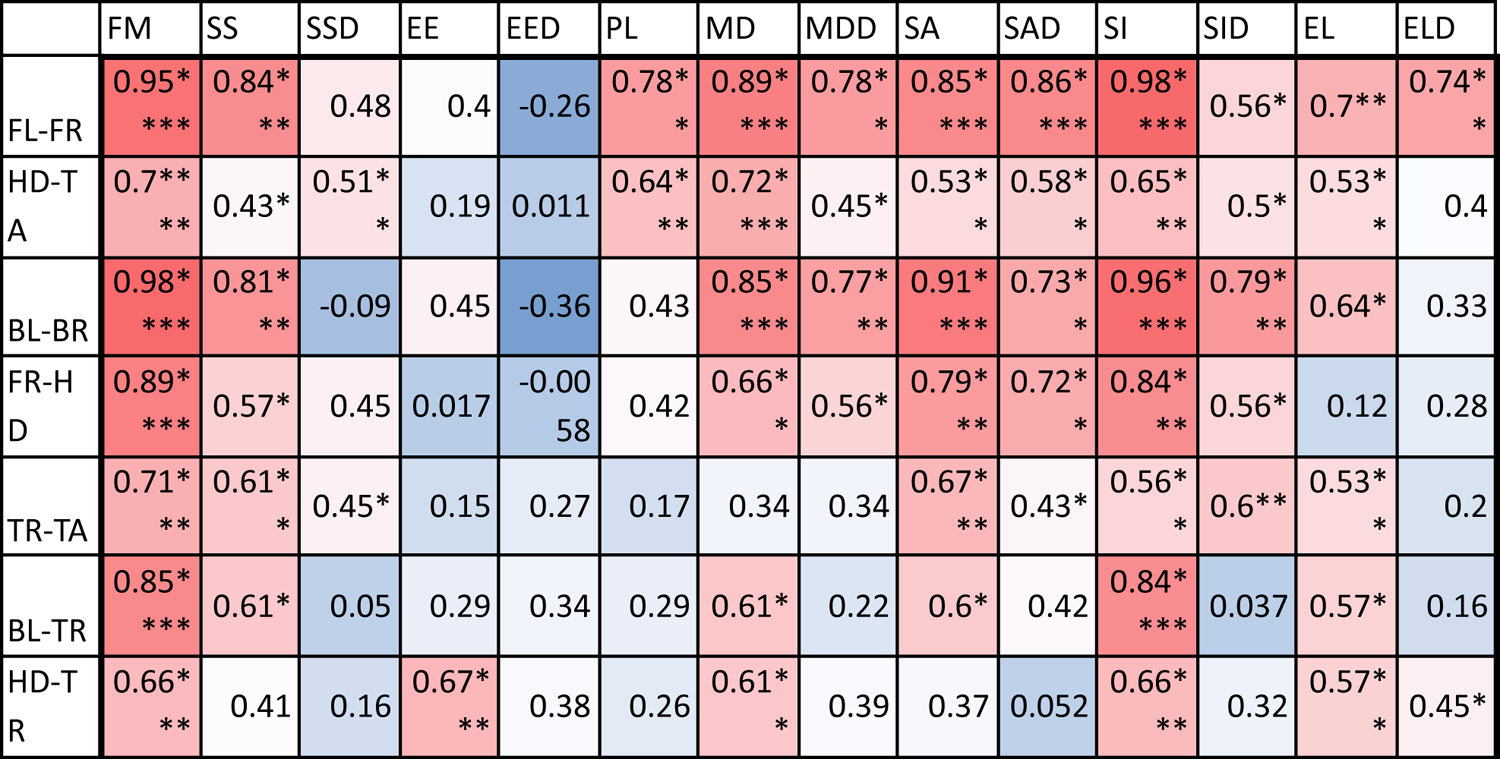
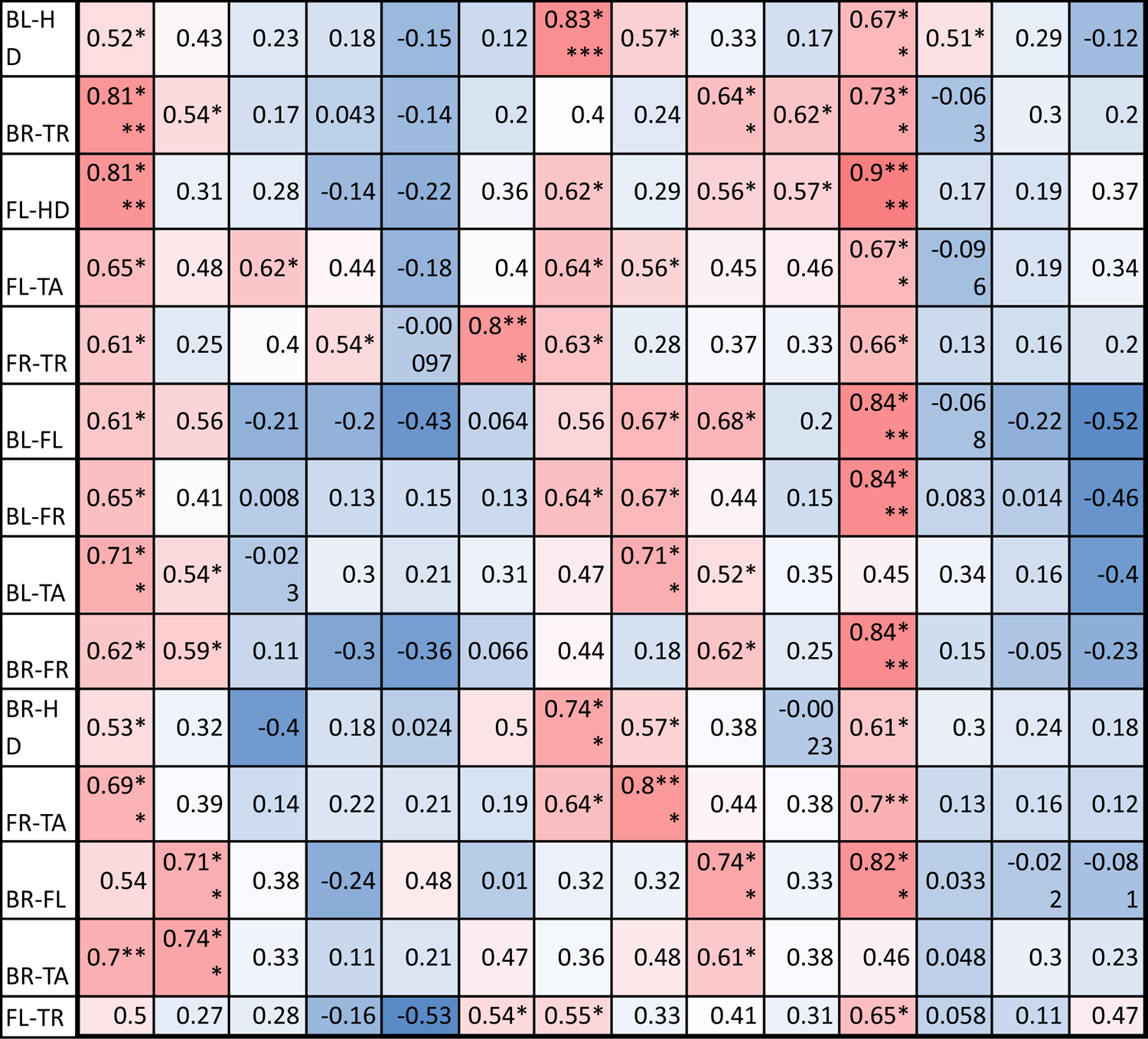
Correlation coefficients of within-individual indices with values color coded from negative (blue) to positive (red). Each row corresponds to a different pair of body parts indicated as in Figure 2. E.g. “BL-HD” denotes comparison of the left back leg and the head of an individual. Columns correspond to the14 indices of pattern characteristics as in Table 3.

Correlation between front and back legs (FL-BL, FL-BR, FR-BL, FR-BR) was much weaker. The trunk pattern was most strongly correlated with the tail pattern (TR-TA; 8 out of 14).

Interestingly, within-individual correlation tends to be weak for indices describing the typical shape of the spot (EE, EL), whereas measures of the relative size of the spots (FM, SS, SA) tend to be highly correlated between body parts.

Correlation coefficients were calculated based on all geckos who had patterns on the corresponding body parts; e.g. the “BL-HD” values are computed over all geckos with patterned back legs and patterned heads. One star (*) indicates p-values less than 0.05, ** p-values less than 0.01, *** p-values less than 0.001, **** p-values less than 0.00001. Rows are sorted by the number of significant correlation coefficients. Data obtained on all the 25 geckos together independently of morphotype. See the Supplementary Material for differences among morphotypes.

We performed a principal component analysis on the 14 indices taken across all the 25 studied geckos and across the different body parts, the results of which are summarized in Table A3 and Figures 4-6. A principal component analysis is a statistical method to convert a set of observations (in our case, among 14 indices that have many overlaps in the information they are describing) into a smaller number of uncorrelated variables called the principal components.

**Figure 4:**
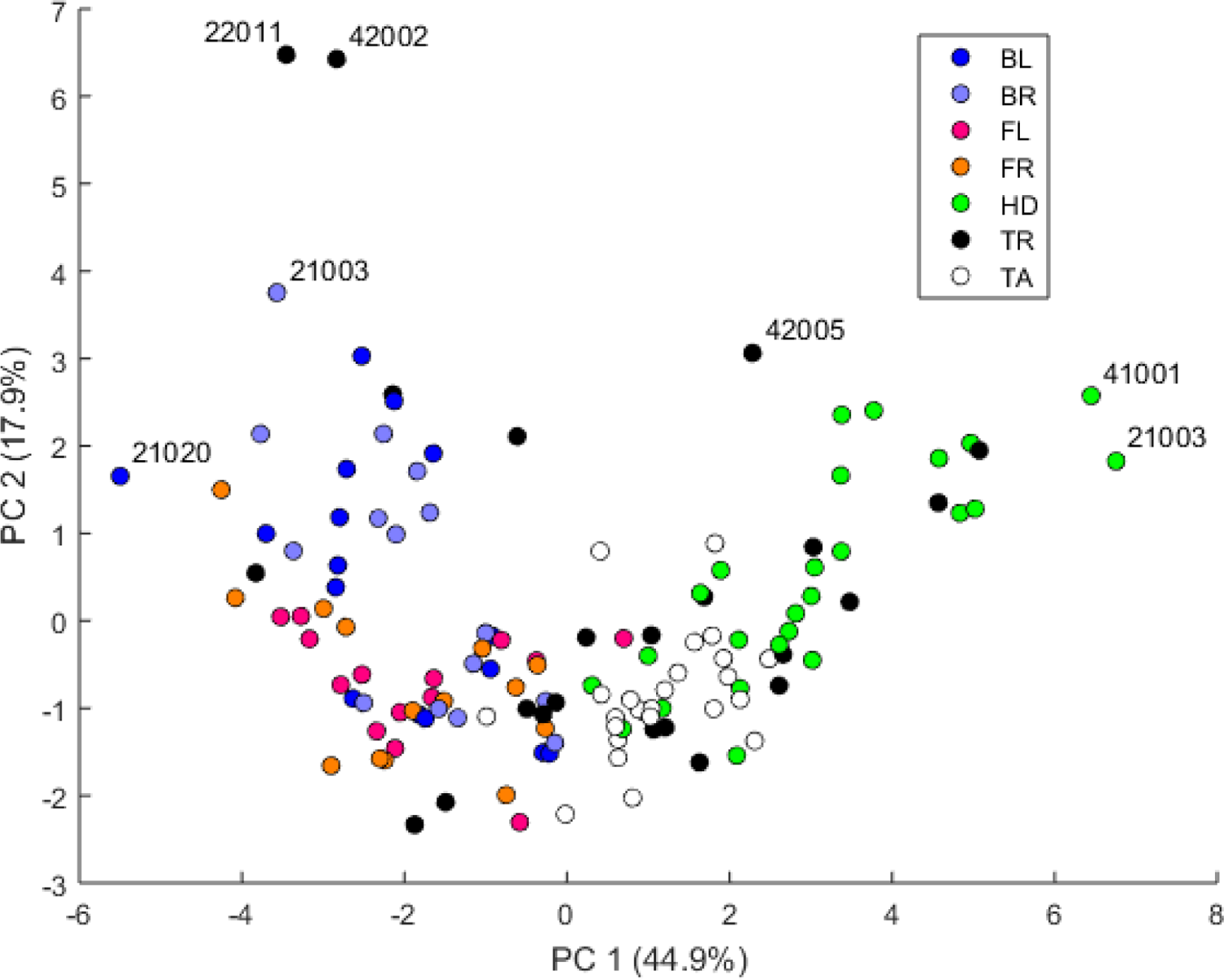
Plot of principal components 1 and 2. Each dot in the plot corresponds to a specific body part of a gecko. Body parts are colored coded and indicated by the same symbol as in Figure 2. A few outliers are labeled via the corresponding gecko id (See Figure A1 in the Appendix for images of all individuals. Note that the two outliers for the trunk patterns in the top left corner correspond to two normal morphs obtained from different sources, suggesting that grouping between these two individuals is not due to them being blood related). Data from all the 25 geckos with pattern in the specific body part indicated.

**Figure 5:**
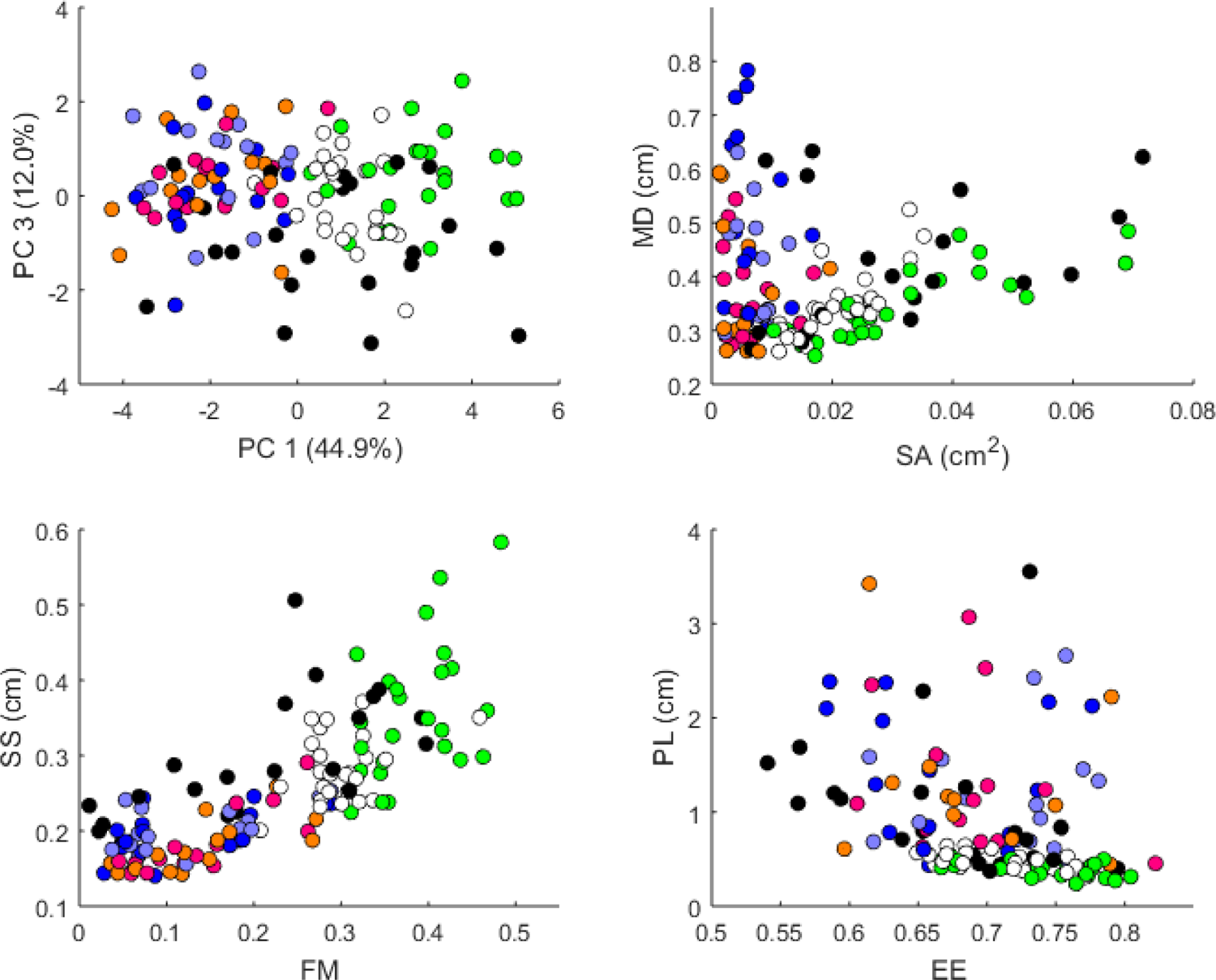
Plots of principal components 1 and 3 (top left); spot area (SA) vs. mean distance between neighboring spots (MD; top right); fractional area (FM) vs. spot diameter (SS; bottom left) and ellipticity (EE) vs. peak length (PL; bottom right). Percentages in parentheses give the fraction of the total standard deviation explained by the given principal component. Body parts are color coded as indicated in Figure 4. Axes bounds are chosen so that in some cases, a few outliers are not shown. Data from all the 25 geckos with pattern in the specific body part indicated.

**Figure 6:**
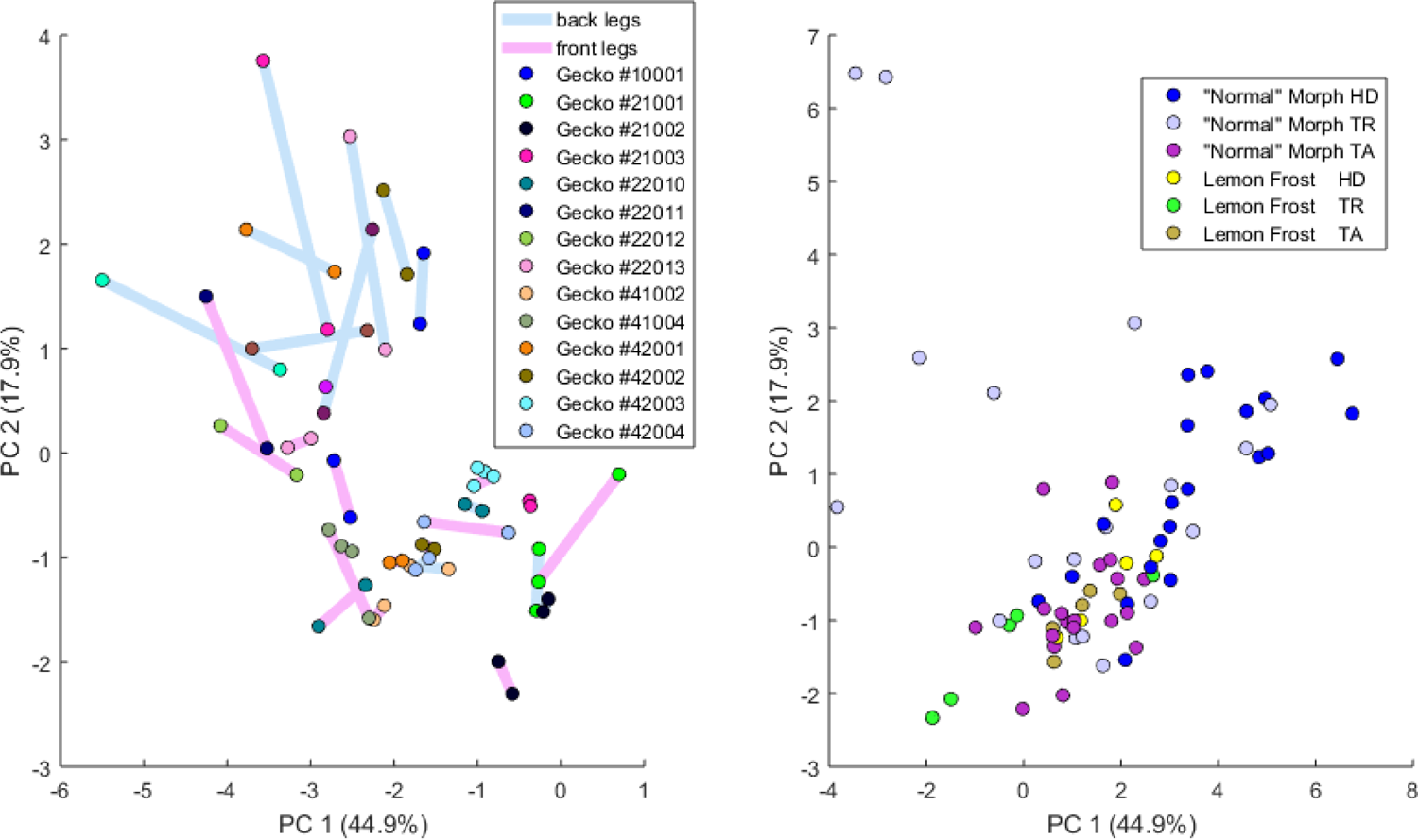
Left: Plot of principal components 1 and 2 of all leg patterns. Geckos are color coded as indicated in the legend and each dot in the plot corresponds to a specific leg of a gecko. The two back legs of the same gecko are connected via a blue line; the two front legs by a purple (‘fuchsia’) line. **Right:** Plot of principal components 1 and 2 of head (HD), trunk (TR) and tail (TA) patterns. Geckos are color coded by morph.

Together, the first seven components account for over 94% of the total variance between the 132 different patterns we analyzed. The first principal component (explaining 45% of the variance) has larger coefficients for those indices that are associated with the average size of the spots, while the second component (18% of the variance) tends to have larger coefficients for those indices associated with the characteristic wavelength (roughly corresponding to the distance among spots) of the pattern. The third principal component (12% of the variance) is strongly associated with EE and EED, which describes how close the individual spots are to circular shapes, and how much they vary in this regard.

Figures 4-6 summarize the PCA data graphically. Together PC1 and PC2 (generally, variation in spot size and separation, respectively) explain nearly two thirds of the pattern variation (62.8%) and provide insight into systematic differences among patterns of the legs, head, tail and trunk. Clusters of leg patterns are well separated and distinct from clusters of the head and tail patterns for plots of PC1 versus PC2 (Figure 4). Since they sort along the PC1 axis (roughly, summarizing measures of spot size), we find that for example the head and tail patterns tend to have larger average spot areas, while leg spots have smaller average spot areas (Figure 5, top right and bottom left). This is true for both the absolute size of the spots (indices SA and SS) as well as the size of the melanistic area as a fraction of total skin area (FM), indicating that leg spots tend to be disproportionately smaller than body spots. When looking at pattern variation only for legs, we observe that the front legs are clustered with lower PC2 values (PC2 is more associated with measures corresponding to the distance among spots) and also variation in the front legs tends to be smaller than variation in back legs (Figure 6, left).

### Within-individual and between-individual distances in pattern space - a measure of the influence of developmental noise in pattern formation

#### i. Pattern distances among pairs of legs

We investigated whether the patterns on each pair of the four legs on each gecko are more similar than the patterns on different geckos. In fact, variation in pattern among the two front legs or the two back legs for each gecko most likely reflects the level of developmental noise in pattern formation on the legs. In Figure 6 (left panel) – where variation reflects mostly differences in the size of the spots and distance among them - a clustering of the legs of individual geckos is not readily visible, suggesting that the contribution of noise generally is more important than the contribution of genetic factors. It should be noted that although a considerable measurement error also contributes to the differences between the legs, random variation due to developmental noise is significantly larger than the measurement error (see Tables A1 and A2). We compared the within-individual differences in pairs of leg patterns to the between-individual differences in the same pairs (Figure 7). We found qualitatively somewhat similar results for both the Mahalanobis distance (gray bars in Figure 7) and the Developmental Noise distance (blue bars in Figure 7) concerning the distance between the two front legs and the distance between the two back legs. In both cases, the within-individual distances were smaller than the between-individual distances. Their ratio was between 0.55 and 0.57 for the Mahalanobis distance and even 0.16-0.19 for the developmental noise metric (a ratio of 1.0 would indicate no difference between the within-individual and the between-individual distances). The differences were highly statistically significant except for the case of back legs and the Mahalanobis distance. The within-individual pair of the two front leg patterns is thus found to be very significantly more similar than two leg patterns from different individuals. (Statistical significance was assessed via nonparametric permutation tests, see the Appendix for more details). Similarly, the within-individual distance between the two back leg patterns is also found to be significantly smaller than the mean between-individual leg distance. For both distances, a front leg and a back leg of the same gecko were closer to each other than a front leg and a back leg patterns randomly taken from two different geckos. However, overall, the mean distance between leg patterns of the same gecko is 64% of the mean distance between patterns on different geckos for the Developmental Noise distance and 91% for the Mahalanobis distance, a not statistically significant difference in the latter case. This further indicates that there is large variation in leg pattern (Figures 4 and 6), and specifically that the variation observed between legs within each individual is only slightly less than the one observed among individuals, consistent with the small within-individual correlation coefficients observed for some indices (Table 5). Our results indicate that within each gecko, two front legs and the two back legs are much more similar to each other than a front and a back leg. This would suggest that the difference between a front and a back leg is not just due to developmental noise. In fact, the results support the hypothesis that the mechanisms - including timing of it - of pattern formation and/or regulation of pattern establishment are distinct for the front and back legs.

**Figure 7:**
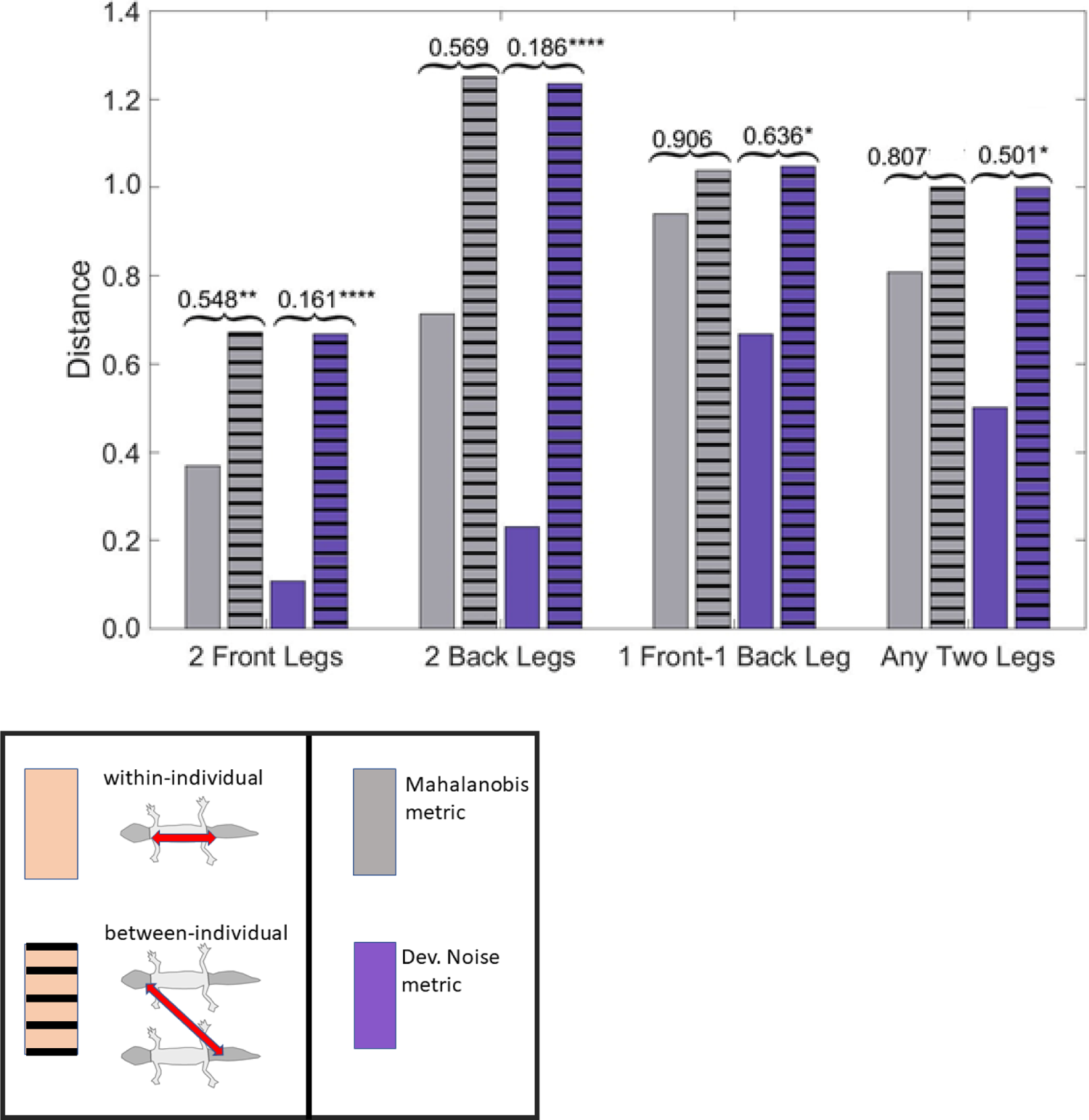
(Squared) distances among leg patterns. For each type of leg comparison, within individual distance (bars without stripes) and between individual distance (bars with stripes) and their ratio (value indicated above each pair of bars) are shown. Gray bars show the Mahalanobis distance, blue bars show the Developmental Noise distance (see inset below the figure). Distance squares are scaled so that the mean between-individual leg distance, i.e. the distance between two leg patterns of different individuals, is 1 in each metric. Stars are based on the p-values for the null hypothesis that the mean within-individual distance is greater or equal to the between-individual distance; equivalently, that the ratio between the two is greater or equal to one. The alternative hypothesis is that the mean within-individual distance is strictly less than the between-individual distance. Data obtained on all the 25 geckos together independently of morphotype. One star (*) indicates p-values less than 0.05, ** p-values less than 0.01, *** p-values less than 0.001, **** p-values less than 0.00001.

#### **ii.** Within-individual and between-individual pattern distances among heads, trunks, tails and legs

We also investigated the distances between the head, trunk, tail, and leg patterns. The results are summarized in Figure 8. The distances are scaled with the same factor as in Figure 7, i.e. the average distance between the patterns of two legs from different geckos is scaled to 1. We compared the within-individual and between-individual distances for all possible pairs of head, trunk, tail and leg patterns.

**Figure 8:**
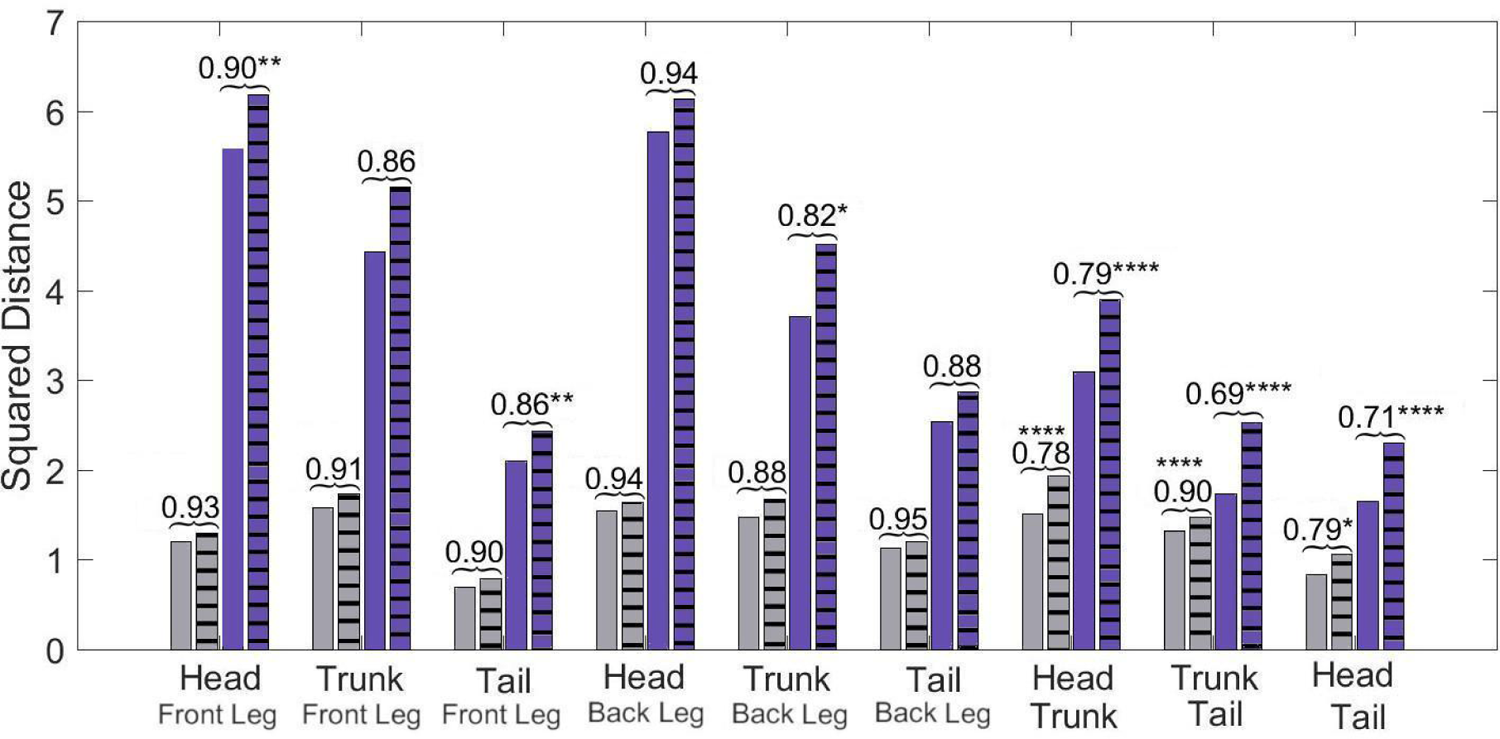
(Squared) distances among head, trunk, tail, and (front) legs. For each type of leg comparison, within individual distance (bars without stripes) and between individual distance (bars with stripes) and their ratio (value indicated above each pair of bars) are shown. Blue bars show the Mahalanobis distance, gray bars show the Developmental Noise distance (see also the legend to Figure 7). Scaling of distances as in Figure 7. Stars are based on the p-values for the null hypothesis that the mean within-individual distance is greater or equal to the between- individual distance; equivalently, that the ratio between the two is greater or equal to one. The alternative hypothesis is that the mean within-individual distance is strictly less than the between-individual distance. One star (*) indicates p-values less than 0.05, ** p-values less than 0.01, *** p-values less than 0.001, **** p-values less than 0.00001. Data obtained on all the 25 geckos together independently of morphotype.

The comparisons of leg patterns with other body parts (legs-head, legs-trunk, legs-tail) yielded that the mean within-individual distance was slightly smaller than the mean between-individual distances for both the Mahalanobis distance and the Developmental Noise distance. However, the differences were relatively small, between 5% and 12% for the Mahalanobis distance and 6% and 18% for the Developmental Noise metric. For the Mahalanobis distance, these differences did not reach statistical significance. In other words, if you take a leg pattern of one individual, the difference between this leg pattern and, say, the head pattern of the same individual is on average not statistically significantly smaller than the difference to the head pattern of a second individual. This is different for the tail, head and trunk patterns. For all three pairs - trunk and tail; head and tail; head and trunk - the within-individual distance is significantly less than the between-individual distance. This means that on average, the patterns of any two body parts of the same animal, e.g. the head and the trunk, is more similar than two patterns from different animals, e.g. the head pattern of one animal and the trunk pattern of another animal. This holds true for both Mahalanobis and Developmental Noise distances.

## Discussion

The goal of this paper was to develop tools and methods to quantitatively study skin pattern variation within and between individuals. We developed a pipeline to collect the data from images of live geckos and then computed various geometric indices to describe characteristics of the patterns. Similar approaches have been taken for giraffe coat patterns (Lee et al., 2018) and salmonid fish skin patterns (Miyazawa et al. 2010); however in our work, not only we captured different aspects of pattern elements and look at variation for each of the elements, but we also used two concepts of distances to quantify the degree of similarity of patterns as whole.

Our method is not only relevant for the analysis of experimental data, but also for the evaluation of mathematical models of skin pattern formation. There are in fact many such models (e.g. Murray, 2002; Cruywagen et al., 1992; Painter, 2001; Cooper et al., 2018; Kondo et al., 2009), encoding various hypothesized mechanisms of pattern formation. It is far from straightforward to rigorously compare the synthetic patterns these models generate to the real skin patterns due to the complexity and irregularity of the patterns. Most authors typically either compare only one or two indices, such as the typical wavelength of the pattern, or just rely on the judgment of human pattern recognition. While this method of comparing patterns “by eye” lacks rigorous quantification, in some ways it is arguably far more developed and sophisticated than current methods of pattern comparison based on lower-dimensional, quantifiable measures. Therefore, our method represents a hybrid approach in which numerous, not necessarily independent measures (the 14 indices used in this work, considered as pattern elements) are chosen at the discretion of a human, based on the pattern variation and characteristics that are perceived as important. The values of these measures are then obtained using automatic methods and mathematical definitions are used to supply suitable weights for the measures to define the distances between patterns. This approach therefore permits to fully depict variation in melanistic patterns within and among individuals and to quantify differences.

## Hierarchy of patterned body parts based on developmental sequence of melanistic patterning

We found an identifiable hierarchy of melanistic patterning head/ tail→ trunk→legs in the studied leopard geckos; for example presence of patterns on the front legs also entails patterns on the trunk, tail and head, but not necessarily on the back legs (Table 1). There is no clear hierarchy between the front legs and the back legs, although for the “normal” geckos in our sample, unpatterned front legs implied unpatterned back legs. These results point to a corresponding order of the establishment of patterns during development: it appears that pattern formation occurs simultaneously in an anterior-posterior and a proximal-distal direction, forming first on the head, then on the trunk, followed by the legs. Patterning of the front legs and the back legs appears to be independent, due to the independent presence of pattern, the low correlation of the pattern indices (Table 5) and the fact that within-individual comparisons of front and back legs yielded very similar results as between-individual comparisons (Figure 7). Pattern formation and establishment on the tail appears to be based on a related mechanism as the head and trunk, as indicated by the similarity of tail and head patterns, as well as tail and trunk patterns. The observed hierarchy of patterning partially follows pigmentation development in this species.

Melanistic pigmentation in the leopard gecko starts to appear around the developmental stage 40 (hatching occurs at stage 42) as a banded pattern on the body and spots on the front legs (but not on the back legs) (Wise et al., 2009). At the beginning of developmental stage 41, the banded pattern is clearly distinct across the body while a spotted pattern occurs on the upper part of the front leg. However, by the end of this stage, pigmentation is occurring across the whole body (Wise et al. 2009). Although the body (head, trunk, and tail) of hatchling and juvenile leopard geckos generally presents a banded pattern, this species undergoes ontogenetic color changes, with adults generally having a spotted pattern (Figure 1) (Landová et al., 2013). Ontogenetic change in color pattern however does not occur for the legs after hatching (see above).

Therefore, head, tail, and trunk follow a process of pattern development and establishment that is different from the one occurring on the legs, with the pattern on the front legs establishing before the one of the back legs (Wise et al. 2009), potentially explaining why absence of pattern is more common in the back legs than the front legs and in the front legs more than in the rest of the body.

## General pattern variation across geckos and body parts

### Variation and correlation among pattern indices

We found large variation and strong correlation among indices related to the amount of melanistic area and the density and size of the spots among the different studied geckos and among the different body parts (Tables 4 and 5). Although variation in spot size across body parts and among individuals may partially be related to size differences, this is less the case for the fraction of melanistic area on the total area. Independently on the observed variation, once spots are formed, their average shape is similar across individuals (Table A4), suggesting a strong constraint on this pattern element (EE), which is also weakly correlated to the other indices (Table 4). While variation in spot size and density has also been observed in other organisms (e.g. Asai et al., 1999; Morgan et al., 2014; Rudh et al., 2007; Balogová and Uhrin, 2015; Druml et al., 2017), less is known about variation in spot shape. Potential genetic or developmental mechanisms may have evolved to ensure maintenance of spot shape and low variability of this trait. On the other hand, other elements of the spotted pattern (e.g., density and size) may be freer to vary in a coordinated way – as suggested by the observed high positive correlation between some indices. Future research could further investigate if low variation in spot shape also occurs in other spotted vertebrates and if it is similarly achieved across organisms. In zebrafish, different alleles of the *leopard* gene result in changes in spot size, density, and connectivity among spots, suggesting that this gene may regulate the synthesis of an activator in a model of reaction-diffusion pattern formation (Asai et al., 1999). Later studies identified the role of *leopard* in regulating interaction among melanophores (or among xanthophores) and in controlling boundary shape for the spots (reviewed in Kondo et al., 2009; Singh and Nüsslein-Volhard, 2015). Similarly, in horses, two genes with different alleles determine the occurrence and amount of melanistic spots on a white colored coat (Druml et al., 2017). The availability of the leopard gecko genome (Xiong et al., 2016), the relative easiness to breed this species, and the existence of CRISPR-Cas9 technology already tested to create mutations in lizards (Rasys et al., 2019) will allow to develop future research to uncover the genetic basis of variation in pattern elements in this species, similarly to what has been done for mammals and other non-mammalian model species.

## Variation and correlation in pattern among body parts

Phenotypic correlation among traits, in this case the correlation of patterns among different body parts of the same individual, may provide information on how these patterns are related developmentally. Phenotypic correlation was investigated in two ways. The first is the standard method of Pearson correlation coefficients for each pair of body parts and for each measurement, summarized in Table 5. The second measure is the ratio of the mean within-individual distance and the mean between-individual distance in pattern space for each pair of body parts, summarized in Figures 7 and 8. Patterns on the legs are statistically almost independent of patterning on the head, trunk and tail. In contrast, the similarity of head, trunk and tail patterns, as well as the similarity of the two front legs and the two back legs for the same animal are statistically significant for both metrics. The similarity of pattern variation observed on the head, trunk and tail suggests that patterning mechanisms are most likely not independent among these body parts, and the same holds for the two front legs and the two back legs (Figures 7 and 8; see also Table 5 for each index separately). However, as melanistic patterns in the legs, and especially the back legs, are more variable and independent in their variation from the rest of the body, this may indicate a different timing or developmental mechanisms of pattern formation and establishment in these body parts. In this sense, the relatively easiness of captive-breeding of this species may provide a unique opportunity into understanding the underlying genetic and developmental processes and mechanisms producing the the observed variation in color pattern in the different body parts (for similar questions, see Cieslak et al., 2011; Druml et al., 2017; Wasik et al., 2014).

### Comparison of within-individual and between-individual differences in leg patterns as a measure of developmental noise

A within-individual comparison of the two front leg patterns yields that the front legs of an individual gecko are significantly more similar than two between-individual front leg patterns. The same holds true for the two back legs. A simple measure of the magnitude of the contribution of developmental noise is given by the ratio of the mean within-individual distance and the mean between-individual distance, which is the amount of variation presumably due to developmental noise alone normalized by the average amount of pattern variation due to all sources (including genetic and environmental). A ratio of 0 would indicate that the legs of individuals show no variation at all within a gecko, meaning that developmental noise plays no part in the establishment of patterns at all. Conversely, a ratio of 1would mean that the variation between leg patterning of the same animal is indistinguishable to the variation between patterning for two different animals. This would indicate that the process of patterning even between geckos would be entirely dominated by random noise. In our data, the contribution of developmental noise to patterning is quite large by this measure with ratios of within-individual distances to between-individual distances between 0.55 and 0.16 or the Mahalanobis and Developmental Noise metrics, respectively, for the front legs and 0.57 and 0.16 for the back legs. While the measurement error contributes to this estimate - it accounts for between 47% and 33% of the within-individual distances (Table A1) - the variation due to developmental noise exceeds this error significantly (Tables A1 and A2). These indices indicate that although variation in color pattern observed within individuals is not produced by developmental noise alone, overall, developmental noise has a very strong influence on this variation. Together with controlled captive-breeding experiments, the combination of mathematical modeling (see section below) and empirical data can be used in the future to further investigate the relative importance of genotype, environment and developmental noise on the variation in color pattern on the different body parts in these animals. Furthermore, our methodological approach can also be applied to other patterned organisms to study similar questions.

### Methodological significance for the analysis of mathematical models

Our method also serves the purpose of establishing a systematic high dimensional quantitative approach to the analysis of synthetic patterns produced by mathematical models of skin pattern formation (see e.g., Murray, 2002; Cruywagen et al., 1992; Painter, 2001; Cooper et al., 2018; Kondo et al., 2009). Indeed, the distance between synthetic patterns and the actual patterns is a quantifiable overall measure of how similar the synthetic patterns produced by such models are to the actual patterns.. More importantly, our method gives a way to quantify the effect of developmental noise on patterns, which in turn can be used to calibrate and test mathematical models of skin pattern formation. More concretely, we can think of the two front leg patterns of an animal as two points in our 14-dimensional pattern space. The two leg patterns are similar, but not identical. These two points represent the variation from a “typical” pattern that corresponds to the environmental and genetic conditions of the particular individual’s development. We postulate the differences between the two legs to be the effect of developmental noise. Indeed, we can conceptualize abstract sets of possible patterns that can be attained under the same combination of environmental and genetic conditions, but with the random contribution of developmental noise. In a previous work, we have called this set a ‘phenotype cloud’ (Kiskowski et al. 2019); see Figure 9 below. In our case, it is a subset of a 14-dimensional pattern space.

**Figure 9:**
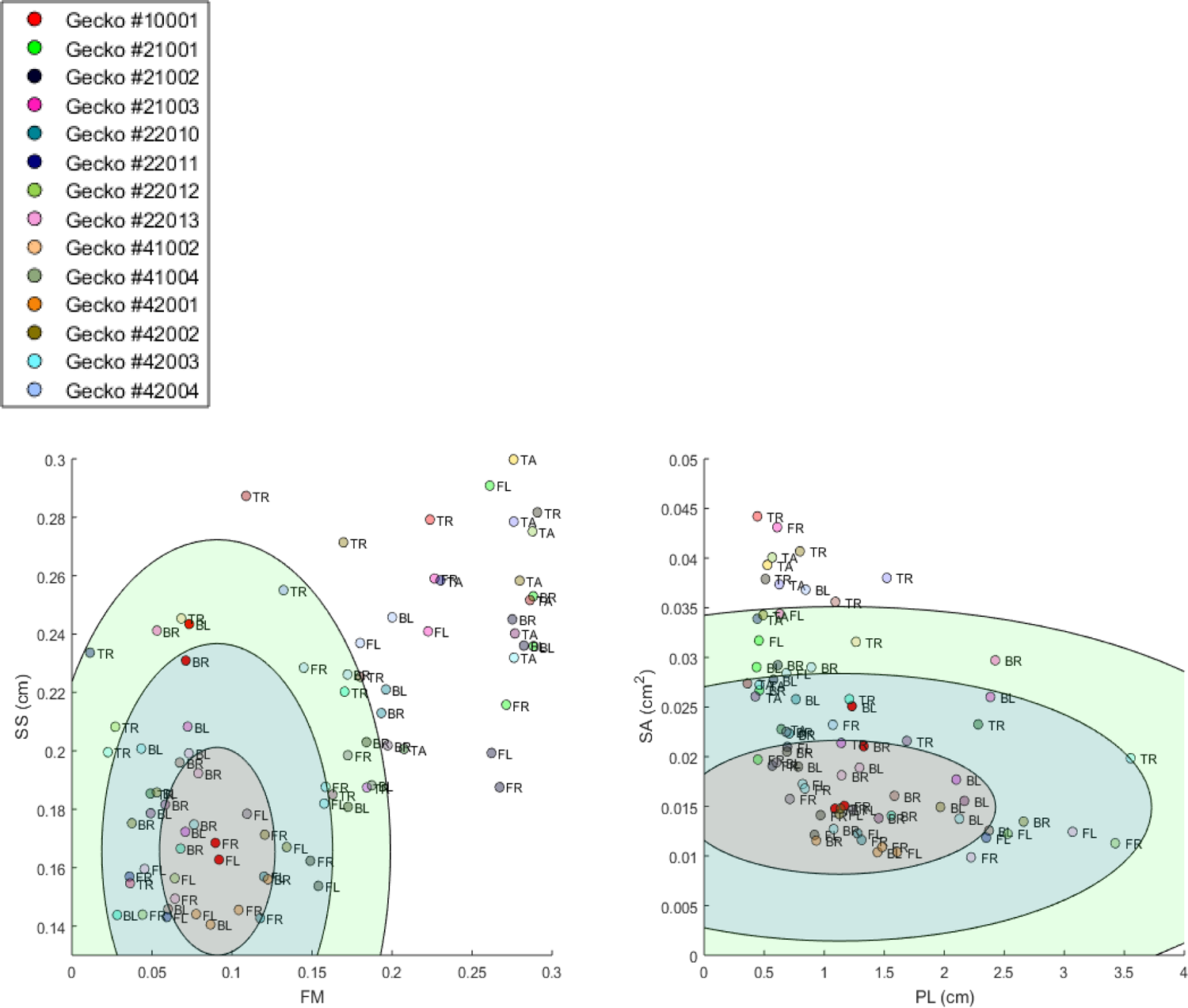
Illustration of the concept of a “phenotype cloud”. The images show the gecko pattern data in FM-SS space (left) and PL-SA space (right). Each gecko corresponds to a unique color and the body part is indicated by a two letter-combination (see Figure 2). The ellipses shown are visualizations of the concept of phenotype clouds of gecko #10001 (indicated in bright red). In fact, these ellipses are projections of certain balls (with respect to the Developmental Noise metric) in 14-dimensional pattern space onto the FM-SS subspace (left) and on the SA-PL subspace (right). These balls are centered at the centroid of the two front leg patterns of gecko #10001 (indicated in bright red). The radius of the innermost ball is equal to the distance of one of the front legs to the centroid in the 14-dimensional pattern space. The other concentric balls have twice and three times this radius. Phenotype clouds can be conceptualized as such balls, e.g. the 95% phenotype cloud is a set that contains randomly generated patterns with the same environmental and genetic conditions and the same level of developmental noise as the front legs of gecko #10001 in 95% of the cases.While it is not possible to generate such phenotype clouds for a single pattern from our data empirically - we only have two data points for each cloud, corresponding to the left and right leg -, it is possible for stochastic mathematical models of pattern formation. The size of the cloud then indicates the contribution of developmental noise to pattern formation - the larger the cloud, the larger the influence of developmental noise.

These phenotype clouds can be interpreted as confidence regions for the phenotypes of individuals with the same genetics and similar environments, extending the ideas of confidence intervals to higher dimensions. One can think for example of a 95% phenotype cloud as an abstract set of patterns that contains a randomly generated pattern (with fixed environmental and genetic conditions, but a random contribution of developmental noise) in 95% of the cases. Since the distribution function for random noise is completely unknown, and we only have two sample points for each instance - the pairs of front legs or the pairs of back legs for each individual-, the actual shapes of these phenotype clouds are unclear. However, in the context of stochastic mathematical models of pattern formation, such model-dependent phenotype clouds can be determined computationally (Kiskowski et al. 2019), which then allows us to test such model prediction against the empirical data.

Thus our approaches can be used to test the validity of mathematical models for skin patterning, and gain insights and formulate prediction on the cellular and genetic mechanisms of pattern formation (e.g., Maini, 2004; Othmer et al., 2009). Besides providing a framework for quantitatively analyzing various aspects of patterns, the concept of the phenotype cloud gives an additional empirical approach to interrogating models.

## Acknowledgements

We are thankful to Gopal Murali, William Allen, and Julien Claude for discussing the correlation of color pattern among distinct body parts in animals during the early phases of writing this article. Julian Claude also provided helpful comments to improve this article. We are thankful to Tony Gamble, Aaron Griffing, and John Scarbrough of Geckoboa for discussion about color pattern and color pattern selection in the pet trade for the leopard gecko and to Matt Vickaryous for discussion about melanistic color pattern formation, especially in regenerated tissues. We also thank Ekkehard Glimm for very helpful discussions about the statistical analysis, in particular for discussions about computations of p-values.

## Author Contributions

TG, MK, and YC thought of and developed the project. All the authors collected the data. TG and MK analyzed the data. TG, MK, and YC wrote the paper.

## Appendix

### Computation of p-values for Figures 7 and 8 and Table A1

We used nonparametric test procedures to compute p-values for Figure 7 and 8 and Table A1. For instance, for the comparison of within-individual and between-individual distances of the front legs,, let µ_within_ denote the mean within-individual square distance between the two front legs, i.e. the mean square distance in pattern space of the two front legs of the *same* animal. Let µ_within_ denote the mean between-individual square distance, i.e. the square distance between front leg patterns of two *different* animals. Fourteen geckos had patterns on both front legs. To compute p-values for the test of the null hypothesis µ_within_ ≥ µ_between_, we generated samples by pairing each of the 14 geckos with another, different gecko in such a way that each of the 14 were chosen exactly once as a ‘partner’. Each such sample corresponds to a permutation of the geckos without fixed point, also called a derangement. For each sample, we chose one of the two front legs of the first partner gecko and one front leg of the second partner gecko. We computed the sample means of the square distances of those two front legs. We repeated this procedure 10,000 times, choosing a randomly selected derangement each time. The reported p-values are the proportions of samples in which the mean sample square between-individual distance was equal to or exceeded the mean square within-individual distance.. Similar permutation approaches were used for the other tests in Figures 7 and 8 and Table A1.

**Figure A1:**
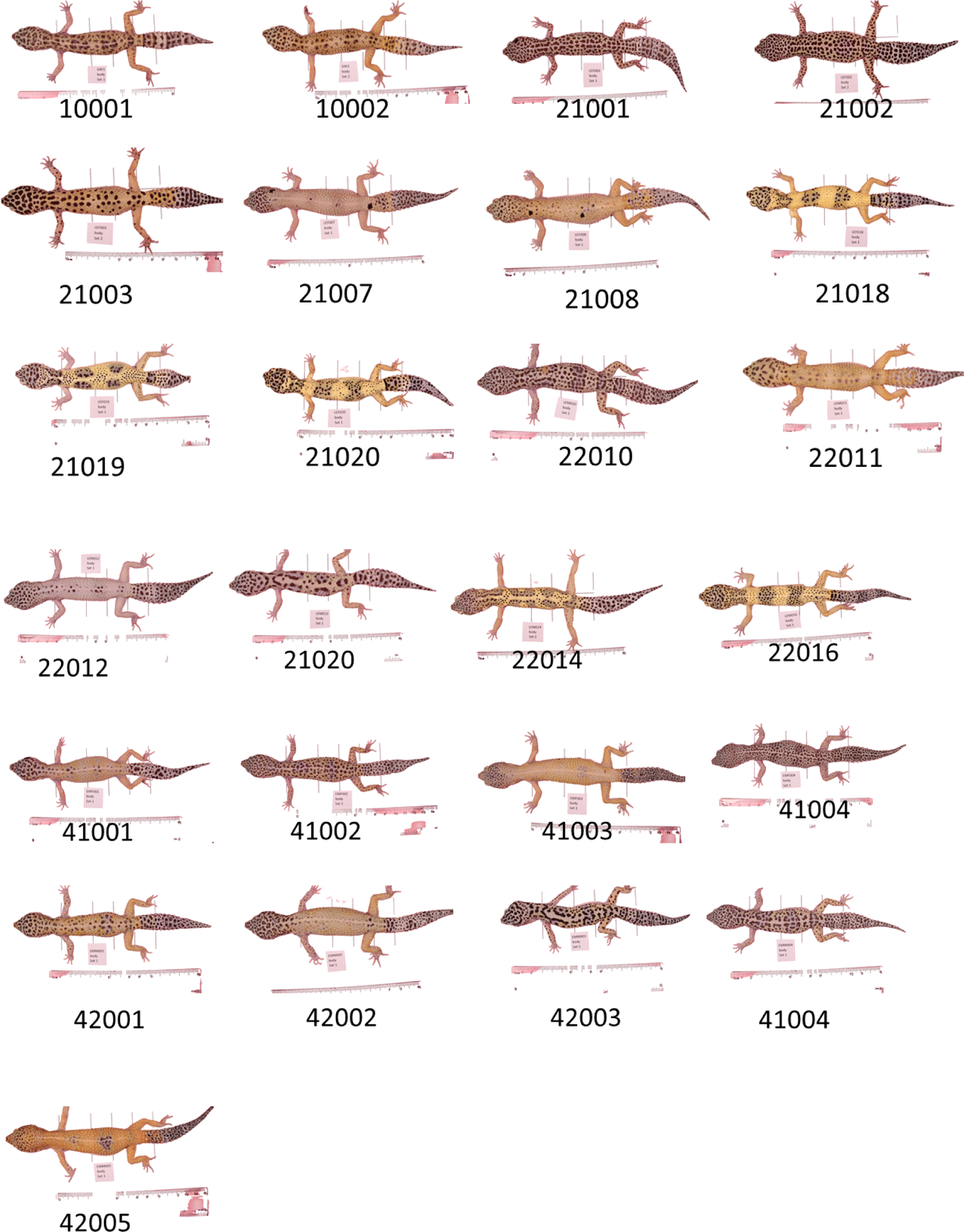
Photos of all geckos included in this study.

**Figure A2:**
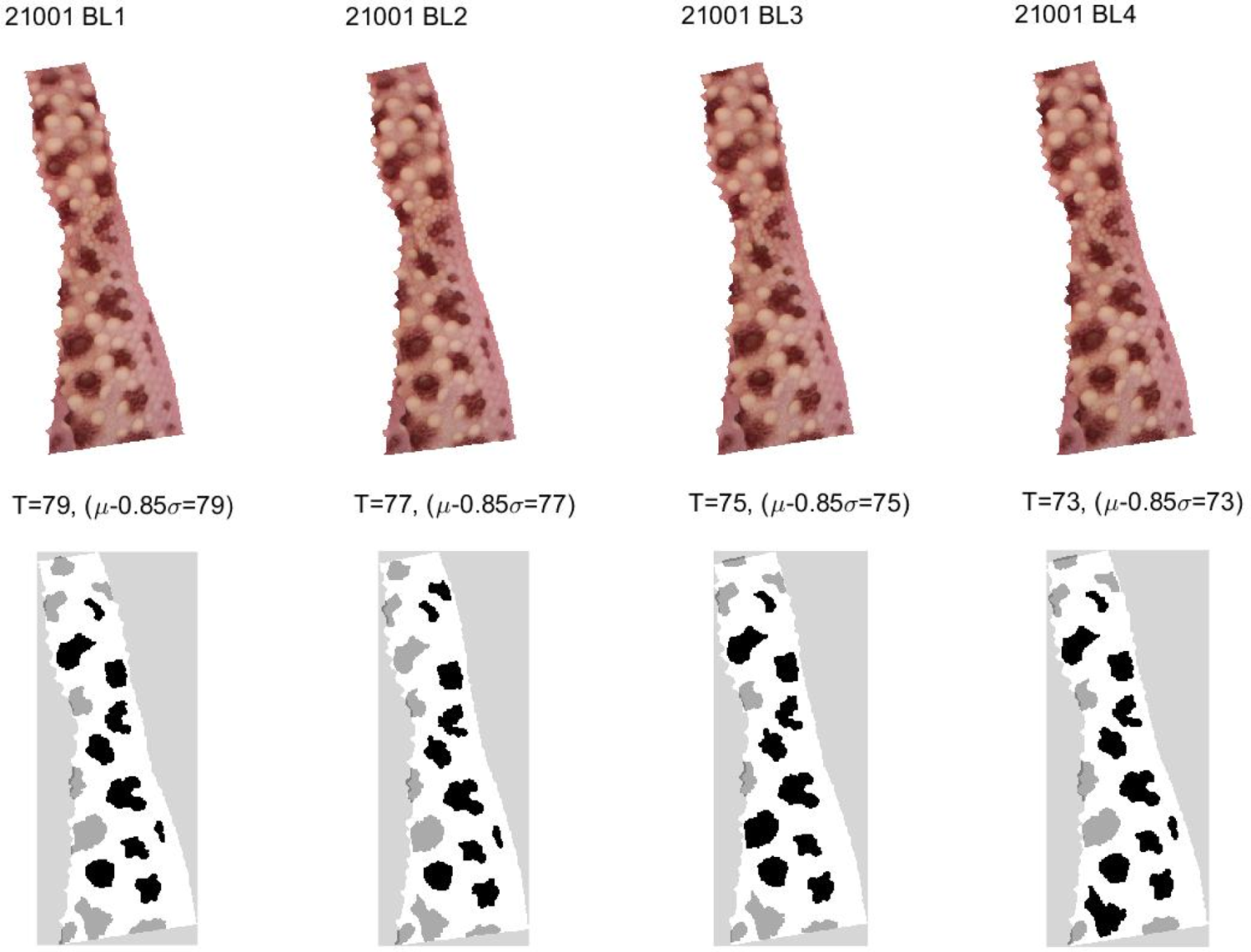
Example of the image binarization: Top: extracted photos of the left back leg of gecko # 21001 for four independent, repeated measurements. Bottom: Binarized image showing spots as detected by the spot detection algorithm for each repeated measurement. Black spots are interior spots, gray ones are boundary spots. The threshold for binarization T is determined by the average pixel value (intensity) µ and the standard deviation of pixel values σfor the observed region of the limb(T = µ − 0. 85σ).

**Figure A3:**
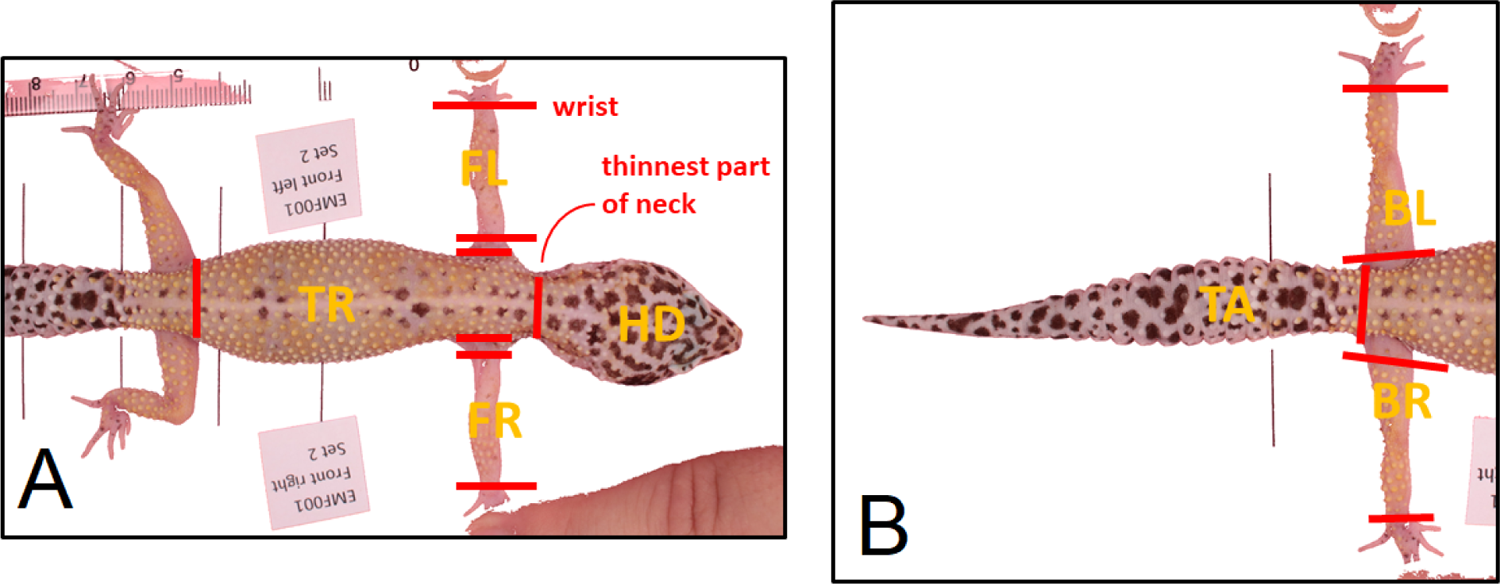
Illustration of body part image extraction. The white background was removed automatically with Matlab by removing pixels in a certain range of RGB values. The red cut were made manually for each image and indicates where boundaries of body parts are.

**Figure S1:**
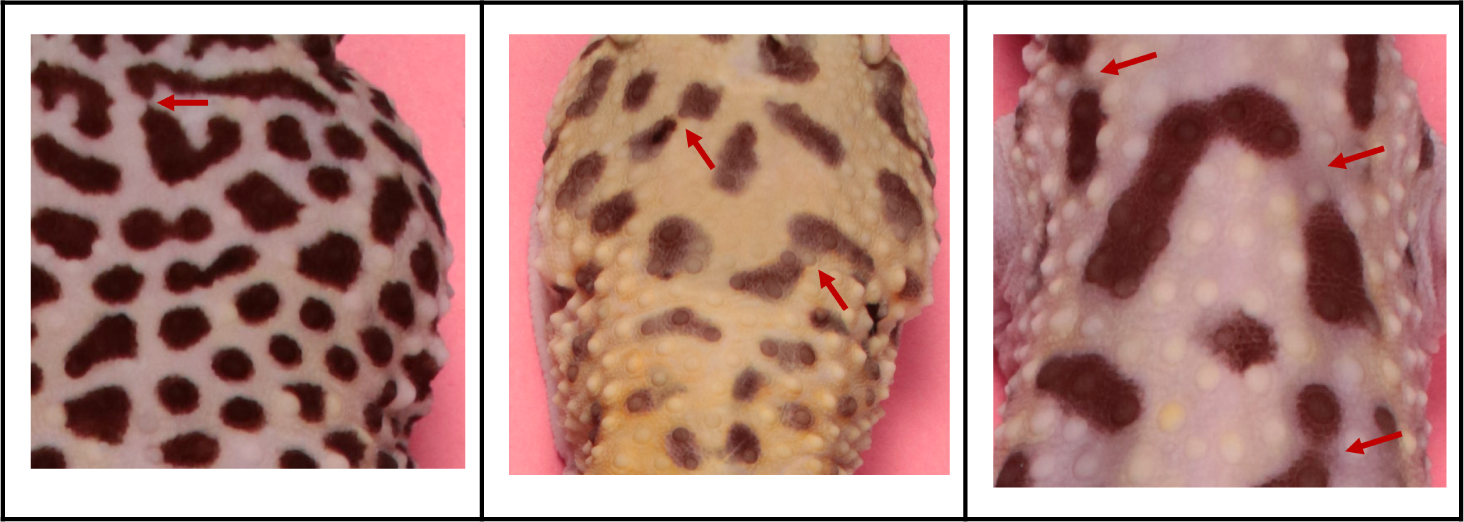
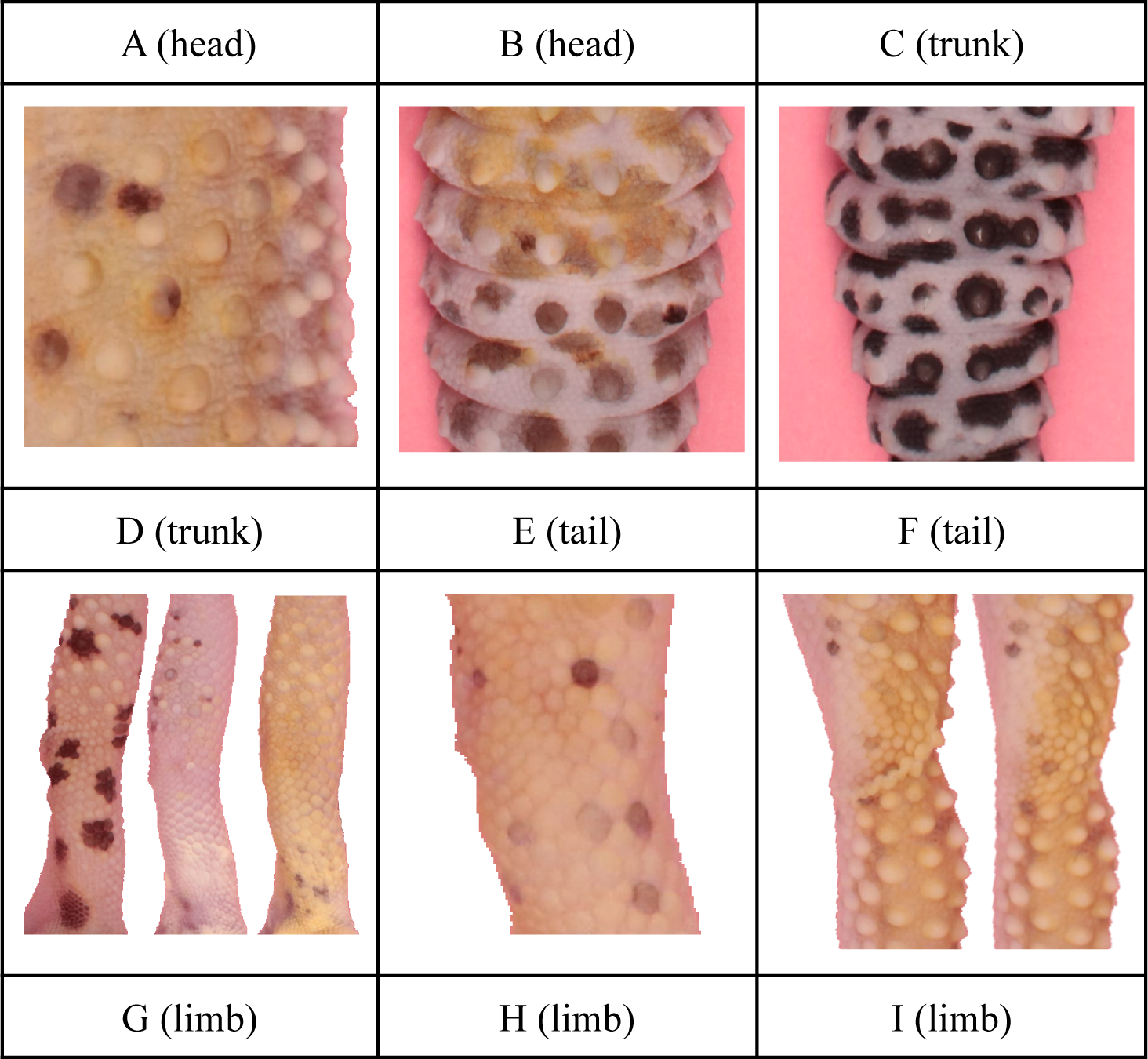
Representative images of patterns found on the four types of body parts; head (A,B) trunk (C, D), tail (E, F) and limb (G, H, I). Head and trunk patterns commonly had very high contrast between spotted and non-spotted pixels (A, C) though not always (B, D) and well-defined spots were often connected by a ‘thread’ with especially light pigmentation (red arrows). The head and trunk patterns often contained spots with a mixture of low and high eccentricity, with spots of both a small, rounded, compact type and a more elongated stripe-like type (A, C). While some patterns could be found with relatively even pigmentation throughout the spots, some patterns had spots with variable pigmentation within and/or between spots (B, D, E, H). Shadows and other types of light effects coincided with the pigmentation pattern due to the rounded contour of the leg (G) and the high protrusion of the tubercules on the trunk (D). Other pattern “defects” included occlusion from wrinkles of the skin that, for example, would appear on some of the replicates but not others (I)

### P-values of correlation coefficients for Tables 4 and 5

The p-values for the correlation coefficients for Tables 4 and 5 were computed with matlab’s *corrcoeff* function (MATLAB R2016b).

**Table A1:**
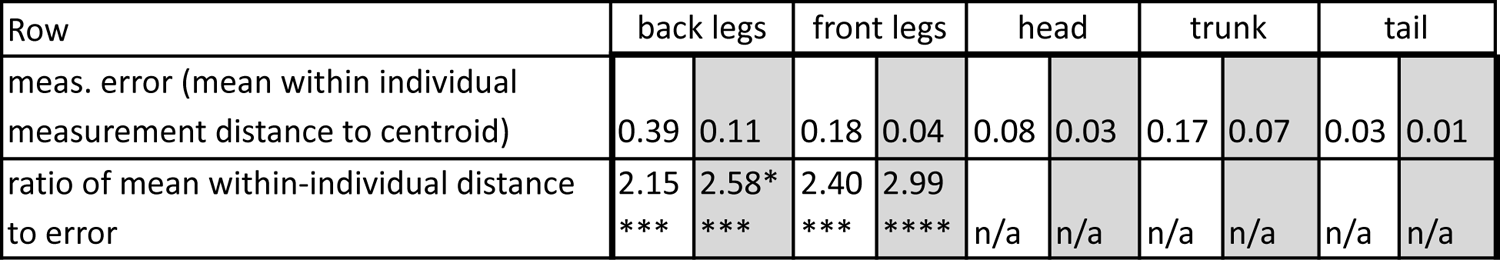
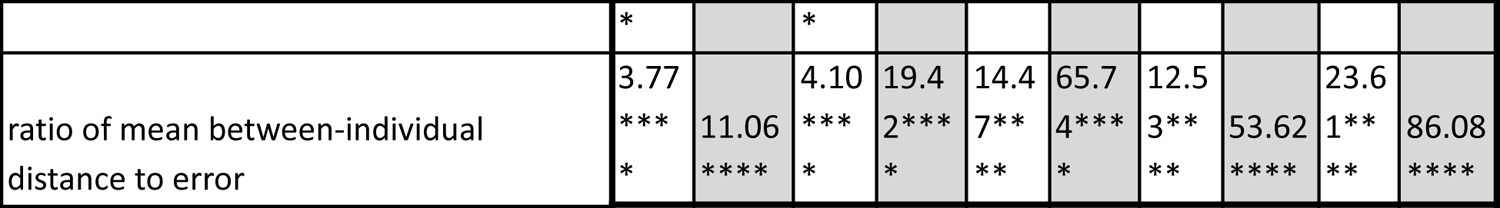
Mean (squared) distances of repeated measurements from the centroid of each body part. White: Mahalanobis distance. Gray: Developmental Noise distance. Distance squares are scaled so that the mean between-individual leg distance, i.e. the distance between two leg patterns of different individuals, is 1, see Figure 7. Given are the ratios of these mean measurement errors to between-individual distances, or, for front legs and back legs, within-individual distances. One asterisk (*) indicates p-values less than 0.05, ** p-values less than 0.01, *** p-values less than 0.001, **** p-values less than 0.00001.

**TableA2:**
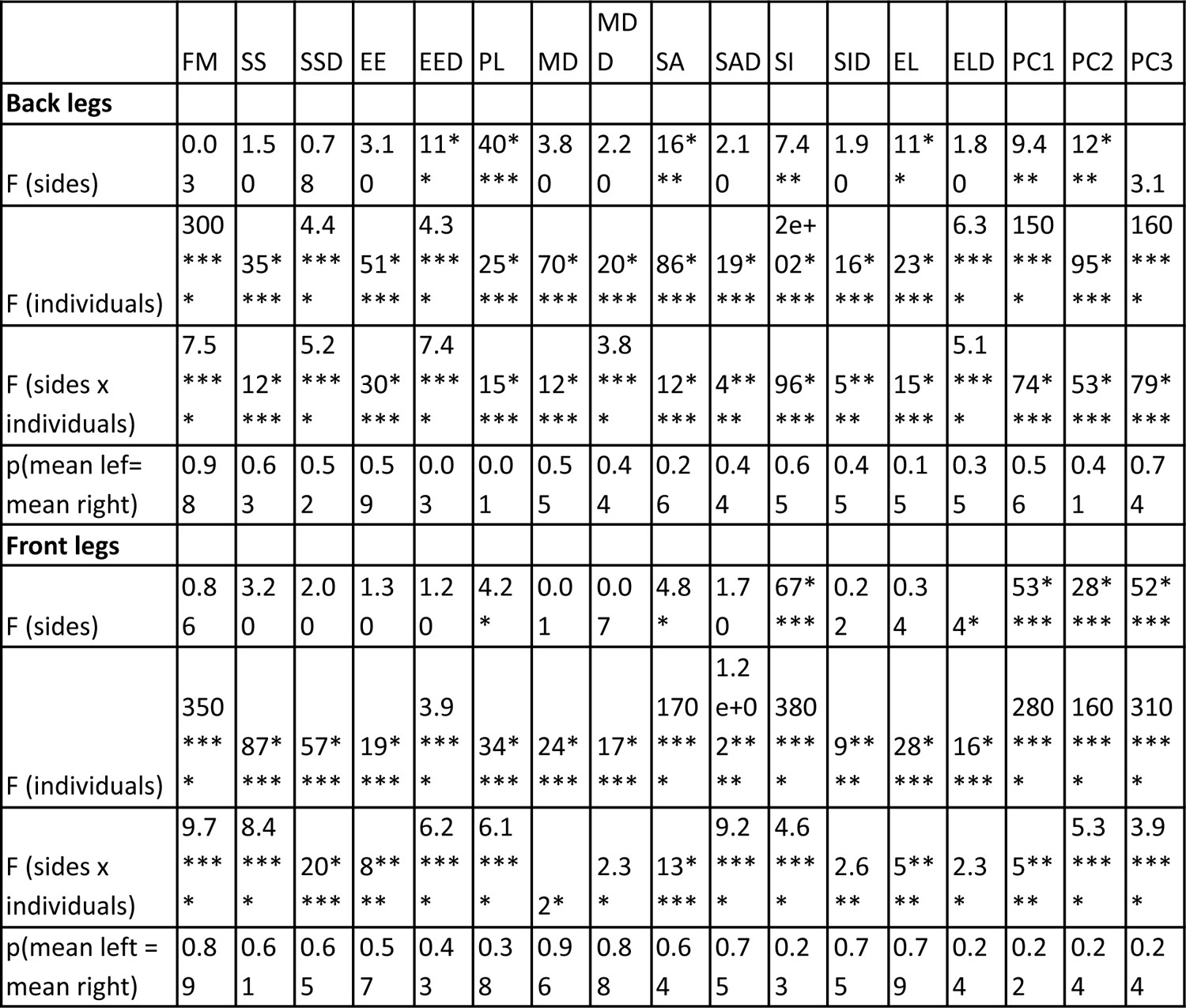
Result of two-way ANOVA test. Given are the F-values (mean square sums relative to the mean square sum of the error) of the factors “individuals”, “sides” and the interaction “sides × individuals”. Also given is the p-value for a t-test of the left versus right means. One asterisk (*) indicates p-values less than 0.05, ** p-values less than 0.01, *** p-values less than 0.001, **** p-values less than 0.00001.

**Table A3:**
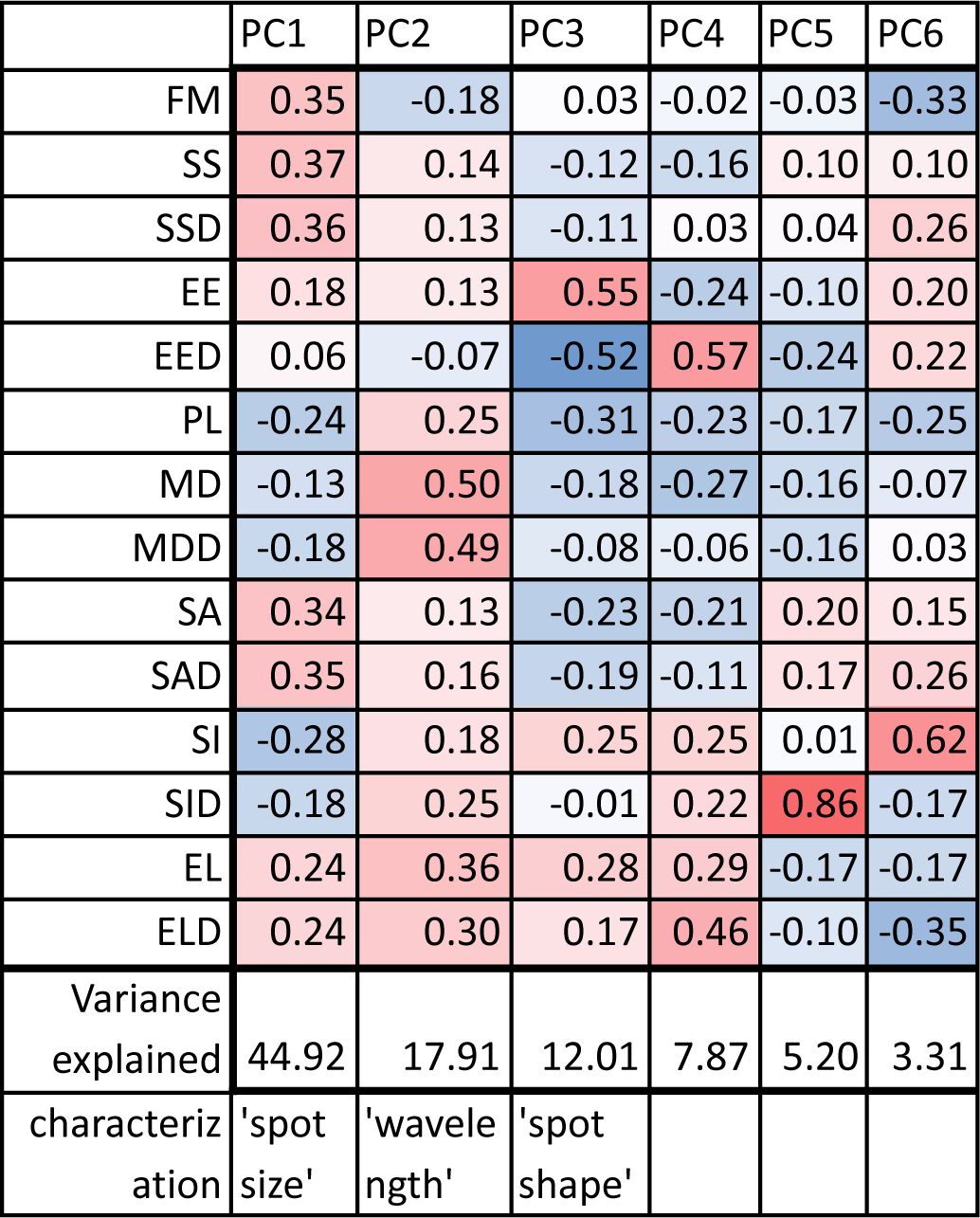
Weights of the first six principal components obtained on the 14 indices from all 132 data points corresponding to different geckos and body parts. Color intensity indicates the magnitude of the coefficients with positive values in red and negatives ones in blue.

|  | PC1 | PC2 | PC3 | PC4 | PC5 | PC6 |
| --- | --- | --- | --- | --- | --- | --- |
| FM | 0.35 | -0.18 | 0.03 | -0.02 | -0.03 | -0.33 |
| SS | 0.37 | 0.14 | -0.12 | -0.16 | 0.10 | 0.10 |
| SSD | 0.36 | 0.13 | -0.11 | 0.03 | 0.04 | 0.26 |
| EE | 0.18 | 0.13 | 0.55 | -0.24 | -0.10 | 0.20 |
| EED | 0.06 | -0.07 | -0.52 | 0.57 | -0.24 | 0.22 |
| PL | -0.24 | 0.25 | -0.31 | -0.23 | -0.17 | -0.25 |
| MD | -0.13 | 0.50 | -0.18 | -0.27 | -0.16 | -0.07 |
| MDD | -0.18 | 0.49 | -0.08 | -0.06 | -0.16 | 0.03 |
| SA | 0.34 | 0.13 | -0.23 | -0.21 | 0.20 | 0.15 |
| SAD | 0.35 | 0.16 | -0.19 | -0.11 | 0.17 | 0.26 |
| SI | -0.28 | 0.18 | 0.25 | 0.25 | 0.01 | 0.62 |
| SID | -0.18 | 0.25 | -0.01 | 0.22 | 0.86 | -0.17 |
| EL | 0.24 | 0.36 | 0.28 | 0.29 | -0.17 | -0.17 |
| ELD | 0.24 | 0.30 | 0.17 | 0.46 | -0.10 | -0.35 |
| Variance explained | 44.92 | 17.91 | 12.01 | 7.87 | 5.20 | 3.31 |
| characterization | 'spot size' | 'wave length' | 'spot shape' |  |  |  |

**Table A4:** Coefficients of variation (ratio of standard deviation and mean) for each of the 14 indices with values color coded from low (blue) to high (red). Rows: values for all body parts across all geckos; last row: values for all data points (geckos and body parts) combined. Symbols for each of the 14 indices are as in Table 3. Data obtained on all 25 geckos together independently of morphotype. See the Supplementary Material for differences among morphotypes.

|  | FM | SS | SSD | EE | EED | PL | MD | MDD | SA | SAD | SI | SID | EL | ELD |
| --- | --- | --- | --- | --- | --- | --- | --- | --- | --- | --- | --- | --- | --- | --- |
| Left back leg | 0.71 | 0.18 | 0.51 | 0.11 | 0.21 | 0.99 | 0.34 | 0.54 | 0.36 | 0.56 | 0.22 | 0.26 | 0.13 | 0.35 |
| Right back leg | 0.64 | 0.15 | 0.48 | 0.08 | 0.37 | 0.53 | 0.49 | 0.63 | 0.32 | 0.44 | 0.21 | 0.23 | 0.12 | 0.43 |
| Left front leg | 0.52 | 0.24 | 0.51 | 0.08 | 0.15 | 0.60 | 0.23 | 0.42 | 0.45 | 0.76 | 0.19 | 0.21 | 0.09 | 0.30 |
| Right front leg | 0.53 | 0.20 | 0.51 | 0.09 | 0.24 | 0.87 | 0.31 | 0.52 | 0.50 | 0.83 | 0.20 | 0.20 | 0.09 | 0.35 |
| Head | 0.13 | 0.25 | 0.31 | 0.06 | 0.18 | 0.20 | 0.19 | 0.28 | 0.50 | 0.49 | 0.20 | 0.23 | 0.13 | 0.28 |
| Trunk | 0.59 | 0.29 | 0.58 | 0.11 | 0.18 | 1.34 | 0.61 | 0.83 | 0.50 | 0.67 | 0.24 | 0.38 | 0.18 | 0.43 |
| Tail | 0.16 | 0.16 | 0.27 | 0.05 | 0.18 | 0.19 | 0.18 | 0.39 | 0.34 | 0.34 | 0.21 | 0.23 | 0.10 | 0.28 |
| All body parts | 0.56 | 0.34 | 0.67 | 0.09 | 0.21 | 1.20 | 0.44 | 0.69 | 0.66 | 0.87 | 0.30 | 0.32 | 0.17 | 0.48 |

### Quantification of measurement error

We took four independent photos of each body part, where the animal was picked up and rearranged for each repetition so that the four measurements would be independent. Although efforts were made to minimize the measurement error by standardizing the overhead lighting and arranging the geckos on a template, measurement error was introduced by slight differences in the rotation and placement, especially for the limbs and tail. While the geckos were sedated, the limbs and tail could be arranged carefully; however, even the muscle tone throughout the limb and tail was variable, affecting the overall contour of the body in subtle ways. Small differences in angles along the contour of the body result in subtle lighting differences that would affect the exact contour of a spot or even whether a spot met the threshold of a spot in borderline cases (see Figure S1, Panels B, D and E for examples of patterns with spots that vary in a continuous way in intensity). Larger differences occur between measurements depending on whether connecting “threads”, which were often of borderline intensity, would meet the requirement for the threshold (several of these connecting threads are shown with red arrows in Figure S1).

In the first approach to characterize the measurement error, we consider body part patterns as points in 14-dimensional phenotype space and use distances between them. We determined the ratio of the mean distance of the four measurements of the same body part of the same animal from their centroid relative to the mean between-individual distances for that body part, or (for front or back legs) the corresponding within-individual differences. This is shown in Table A1 by means of the Mahalanobis distance and the Developmental Noise distance. In all cases, the mean within- or between-individual distances were significantly greater than the mean distance due to the measurement error, with factors varying from 2.2 (back leg, within-individual distance relative to measurement error, Mahalanobis distance) to 86.1 (tail, between-individual distance relative to measurement error, Developmental Noise distance). Table A1 also makes it possible to compare the absolute measurement errors of the different body parts (top row). The back legs had the largest average measurement error, 2-3 times that of the front legs. The front legs had similar measurement errors as the trunk, with much smaller errors for the head and tails.

A two-way ANOVA test where the two factors are “sides” (S; fixed) and “individuals” (I; random) (Palmer and Strobeck, 1986; Merila and Biorklund, 1995; Breuker et al., 2006) was performed separately for both pairs of front legs and pairs of back legs for each of the 14 indices. The data sets were all individuals with patterns on both front legs, or on both back legs, respectively, regardless of morph. The ratio F = MS(S×I)/MS(E), where MS(E) is the mean sum of the squares for the error, can then be used as a measure of the relative size of the measurement error. Results are summarized in Table A2. For each index, an F-test yielded that nondirectional asymmetry is making a significant contribution to the variation observed relative to measurement error. The F-values ranged between 2.3 and 96, with a median value of 6.8, meaning that the measurement error made up between 43% and 1% of the observed variation with the median value corresponding to about 15%. We also tested whether directional asymmetry was present via a two-sided t-test. Except for the indices PL and EED for the back legs, this was not the case (note that the two p-values below 0.05 were 0.03 and 0.01, so given that we performed 28 tests of significance, this is fairly weak evidence for the existence of directional asymmetry even in these two incidences).

## Supplementary Material

### Origin of geckos

Different morphotypes in the leopard gecko are obtained through controlled captive breeding aimed at selecting for certain characters related to body and/or eye color, spots vs. stripes and loss of pattern; an introduction to the different well established morphotypes and a general description of the “normal” morphotype vs. other more or less “pure” morphotypes can be found on several web sites of commercial gecko dealers, see e.g. Sykes (2004) and GeckoBoa Reptiles (2008).

All the geckos used in this study are of captive-bred origin and were obtained from different sources (see Table 1 for details) to ensure variation among individuals in the amount of melanistic patterns that exist for captive bred animals. No animal used in this study was purchased or obtained for their pattern, as we only requested geckos with a “normal” melanistic morphotype, which includes all geckos with a melanistic pattern on a yellowish or brownish background. While geckos with “normal” morph are more similar to the “wild type” pattern of this species - and some of the captive bred individuals may have some wild type in their blood line - this morph is generally very variable in terms of melanistic pattern (e.g., Figure A1 in the Appendix) and individuals are bred without specifically selecting for one or more characters in contrast to what happens with the “pure” and different morphs that can be created in this species (albino, tremper albino, melanistic, lemon frost, etc.). The advantage of using this approach - randomly selecting geckos with a “normal” morph obtained through captive-breeding or colleagues - is that by doing so we ensure having a very variable sampling of geckos for this general “normal” morph, as different people obtain them through breeding distinct parental individuals as long as they have the general “normal” pattern apparence.

### Procedure for image acquisition

Geckos were anaesthetized using an open drop technique with cotton balls embedded with isoflurane. Anaesthetization of the geckos ensured that the animals were asleep while we took pictures for all the four datasets for each animal. Pictures were taken for the entire body of the animal - including the head and the tail if the entire animal fit into the frame - and for the two sets of legs (front and back) separately. Legs were held stretched out by two of us (NM and YC) and placed at the same angle from the body across picture sets. Legs were always positioned on top of the horizontal lines printed on the colored paper, while the body of the gecko was placed on the vertical line. Depending on the entire length of the animal, pictures of the tail were obtained together or separately from the picture of the main body. ID cards with information about the picture set were placed in the frame of all photos for use in identification during processing of images. A ruler was also placed in the frame of all the photos as a size reference. Photos were taken using a photography stand with the camera positioned directly at 12 cm above the gecko. Lighting was provided by the overhead lights in the room and by two additional lamps that were attached to the stand and directed towards the geckos. We used a Canon EOS Rebel T6i camera with focal length of 18mm exposure time of 1/80, aperture of f/8, ISO-400, and auto-focus on. Settings were consistent across all photos and picture sets. To obtain pictures for different body parts (e.g., body vs legs), the animal was moved and then replaced every time for each picture set. We first obtained all the four sets for an animal before obtaining pictures for a new individual.

### Description of patterns on different body part

Each of the four body parts (limb, trunk, head, and tail) presented different challenges so that the criteria for identifying melanistic spots were different for each type of body part. For example, the legs of the gecko were rounded and topographically complex with sharp angles due to the bones of the leg (see Figure S1, Panel G), whereas the head, trunk and tail were relatively flat.

Thus the legs were unevenly illuminated and the algorithm for identifying the pattern of the legs included steps to brighten shadowed regions and darken bright regions. Shadows and glare were identified as contiguous areas of pixels that were darker or brighter than the mean pixel intensity at a spatial scale larger than the spatial scale of the spots. This method was effective since the melanistic spots on the legs were consistently smaller than the spatial scale of the leg contouring for all the gecko morphs. Spots identified by the algorithm almost always coincided with melanistic spots identified via eye inspection; only in a very few cases did the algorithm identify what looked like shadows as spots or vice versa (a comparison of photos and algorithm-identified spots for all patterns is added in the Supplementary Material). For the purposes of objectivity, consistency and reproducibility, the results of the algorithm were always used, even when there was a discrepancy between the algorithm and the pattern identified by eye. The trunk presented the challenge that dark crescent-shaped shadows of tubercules were difficult to distinguish from melanistic spots (see Figure S1, panel D). We found that these shadows were darker than melanistic spots in the blue channel and a satisfactory fraction of these shadows could be distinguished from the melanistic pattern by imposing additional blue channel criteria. Melanistic patterns varied considerably in their relative darkness, from gecko to gecko, body part to body part and even within a regional pattern (see Figure S1, Panels B, D, E and H). A qualifying melanistic pattern was identified by selecting pixels for each image that were sufficiently dark relative to the average pixel brightness of that image. The relative amount of darkness that was required to meet the threshold (see threshold section in “Methods”) depended on the body part and the threshold rule was applied after the image was adjusted for shadows and glares. A lower threshold was chosen for the head because the fraction of melanistic pattern area on the head was typically much higher (for example, see Figure S1, Panel A) so that the average pixel intensity was closer to the intensity of the melanistic spots. This lower threshold was effective for the head patterning, whereas it would not work effectively for the identifying the patterning of the other body parts, since the head patterning had a relatively high contrast between melanistic and non-melanistic regions.

### Identification of interior and edge spots

Due to the fact that pictures were taken from above the geckos and due to the rounded shape of the gecko body, especially for the limbs, spots at the edges of the region were occasionally partially cut off from view. We call such spots whose boundary intersects with the edge of the body region “edge spots”. Spots whose boundary lies entirely within the region of the body in the image are called “interior spots” (see Figure A2 in the Appendix for an example). Some of our measures should be computed with the full contour of the spot, but are not affected if edge spots are removed, while other measures do not require the full contour of the spot, but are affected if edge spots are removed. For example, the fractional melanistic area of spots depends only on the total melanistic area, rather than the spot size and shapes. Other measures such as the average size and eccentricity of spots require the entire contour of a spot but do not depend on the total number of spots. For this reason, some measures were computed for the entire spot pattern, including both interior and edge spots, and some measures were computed only for the interior spots that were not cut off by the edge of the region (see Table 3).

### Differences between “lemon frost” and “normal” morphs

Our data set contained 20 geckos of the “normal” and five of the “lemon frost” morphotypes. Although this sample size is relatively small for the “lemon frost” and as such, the analysis of this section should be regarded as preliminary, a clear qualitative difference in the patterns of “normal” and “lemon frost” morphotypes can clearly be detected by eye, so that it is maybe not so surprising that we still obtain several statistically significant differences even with these small samples. The difference between the “lemon frost” morph and the “normal” is illustrated in Figure 6 (right panel), where we show the first two principal components of the head, trunk and tail patterns of the two morphs. For both the head and the trunk patterns, but not the tail patterns, some clustering is observable with lemon frost geckos tending to have smaller values for both principal components. This corresponds to smaller spots arranged in patterns with a shorter wavelength. In fact, when we investigated the differences between the “normal” and the “lemon frost” morphs for the head and trunk patterns (TableS1) we found that these differences are statistically significant for the first two principal components of the head patterns and the second principal component for the trunk. For individual indices, the differences are statistically significant in a few cases, most notably those that are associated with the shape of the spots (EE and EL). In both cases, the “lemon frost” spots are closer to circular than the “normal” spots. In contrast to this, the differences in tail patterns were not statistically significant in any of the indices or principal components; the smallest p-value of all indices was 0.24. We also investigated whether one of the morphs showed more variation in traits than the other (Table S1). With some exceptions, the sample standard deviations of the “lemon frost” morph are smaller than those of the “normal” morph, meaning that the “lemon frost” morph tends to have smaller variability than the “normal” one. This is particularly true for the head and trunk patterns, but there is again relatively little difference between the variances of the tail. Most of these differences are not statistically significant due to the small sample size with only 5 “lemon frost” individuals, but for the trunk, both MD and PL are significantly smaller (along with 5 other indices or principal components), and for head, MD is significantly smaller as well (with 4 other variables also). Both MD and PL are measures of the characteristic wavelength of the pattern, i.e. the typical distance between spots. Thus our results indicate that the “lemon frost” morph has less variability in this measure than the “normal” one for the head and trunk.

**Table S1:**
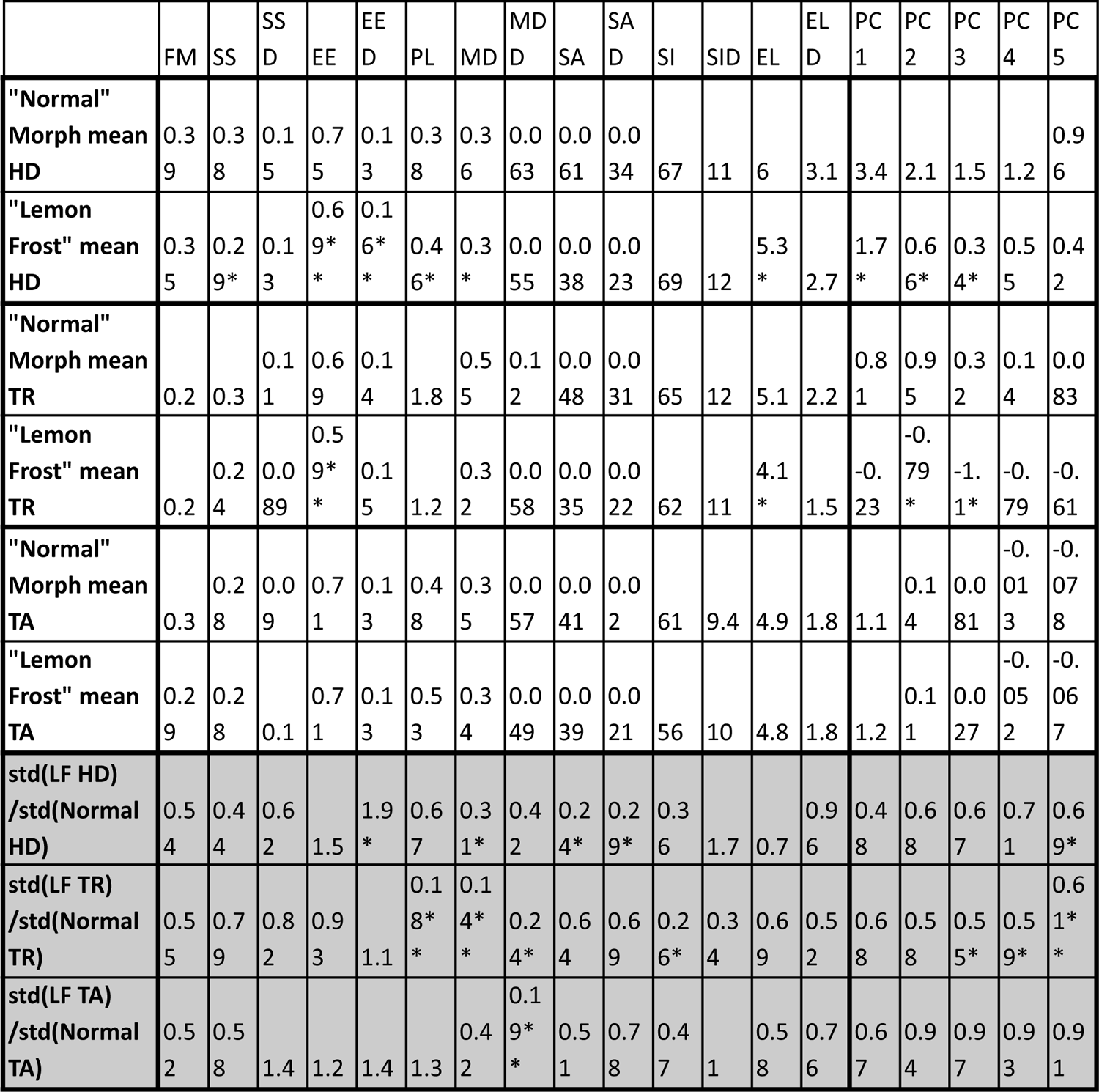
First four rows: Mean values for all 14 indices and the first 5 principal components for the head, trunk and tail patterns of the “normal” and the “lemon frost” morphs. Stars in rows 2, 4 and 6 indicate the results of tests of significance of the hypothesis that the two mean are equal (two-sample t −test). Fifth and sixth row: Ratios of the sample standard deviations of the “lemon frost” morphs and the “normal” morphs. Stars indicate the results of tests of significance of the hypothesis that the two variances are equal (two-sample *F*-test for equal variances). One star (*) indicates p-values less than 0.05, ** p-values less than 0.01, *** p-values less than 0.001, **** p-values less than 0.00001.

### Differences between sexes

Finally, we also investigated the differences between female and male patterning, restricting the analysis to the head patterns of individuals with “normal” morphs. We concentrated on the head patterns because these had the smallest measurement error. None of our indices showed any significant differences (Table S2), consistent with the hypothesis that there is no significant difference between female and male melanistic patterns in captive bred individuals.

**Table S2:**
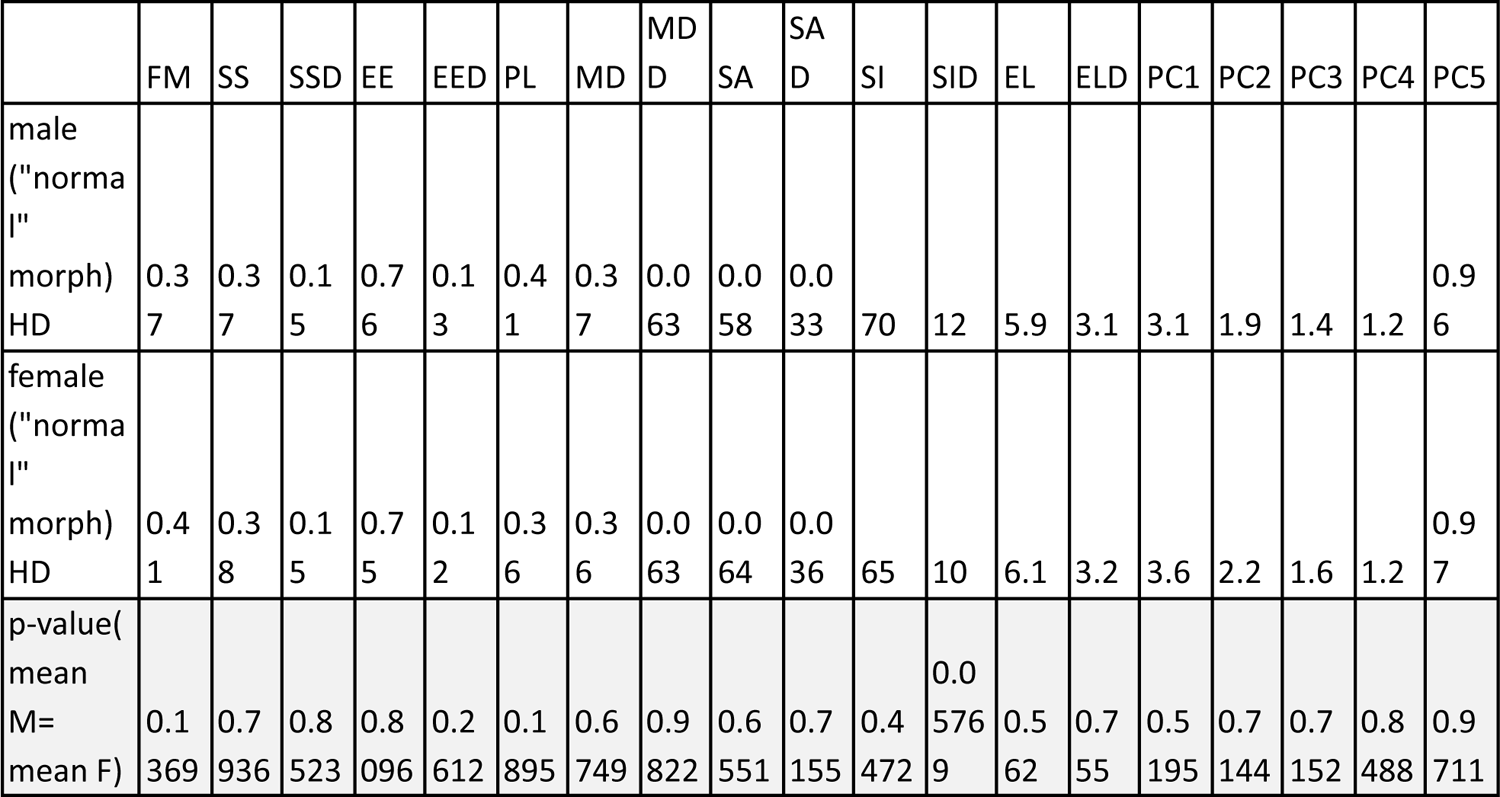
Mean values for all 14 indices and the first 5 principal components for the head patterns of male and female “normal” morph geckos. The third row gives p-values for tests of significance of the hypothesis that the two mean are equal (two-sample t −test). Note that all p-values are well above 0.05, meaning that we cannot reject the hypothesis of equal means for any of the indices or principal components.

Supplementary Material: Data set of all body part images with binarizations and matlab code for image analysis and statistical analysis (available after manuscript acceptance)

